# SARS-CoV-2 proteases cleave IRF3 and critical modulators of inflammatory pathways (NLRP12 and TAB1): implications for disease presentation across species and the search for reservoir hosts

**DOI:** 10.1101/2020.06.05.135699

**Authors:** Mehdi Moustaqil, Emma Ollivier, Hsin-Ping Chiu, Sarah Van Tol, Paulina Rudolffi-Soto, Christian Stevens, Akshay Bhumkar, Dominic J.B. Hunter, Alex Freiberg, David Jacques, Benhur Lee, Emma Sierecki, Yann Gambin

## Abstract

The genome of SARS-CoV-2 (SARS2) encodes for two viral proteases (NSP3/ papain-like protease and NSP5/ 3C-like protease or major protease) that are responsible for cleaving viral polyproteins for successful replication. NSP3 and NSP5 of SARS-CoV (SARS1) are known interferon antagonists. Here, we examined whether the protease function of SARS2 NSP3 and NSP5 target proteins involved in the host innate immune response. We designed a fluorescent based cleavage assay to rapidly screen the protease activity of NSP3 and NSP5 on a library of 71 human innate immune proteins (HIIPs), covering most pathways involved in human innate immunity. By expressing each of these HIIPs with a genetically encoded fluorophore in a cell-free system and titrating in the recombinant protease domain of NSP3 or NSP5, we could readily detect cleavage of cognate HIIPs on SDS-page gels. We identified 3 proteins that were specifically and selectively cleaved by NSP3 or NSP5: IRF-3, and NLRP12 and TAB1, respectively. Direct cleavage of IRF3 by NSP3 could explain the blunted Type-I IFN response seen during SARS-CoV-2 infections while NSP5 mediated cleavage of NLRP12 and TAB1 point to a molecular mechanism for enhanced production of IL-6 and inflammatory response observed in COVID-19 patients. Surprisingly, both NLRP12 and TAB1 have each two distinct cleavage sites. We demonstrate that in mice, the second cleavage site of NLRP12 is absent. We pushed this comparative alignment of IRF-3 and NLRP12 homologs and show that the lack or presence of cognate cleavage motifs in IRF-3 and NLRP12 could contribute to the presentation of disease in cats and tigers, for example. Our findings provide an explanatory framework for in-depth studies into the pathophysiology of COVID-19 and should facilitate the search or development of more effective animal models for severe COVID-19. Finally, we discovered that one particular species of bats, David’s Myotis, possesses the five cleavage sites found in humans for NLRP12, TAB1 and IRF3. These bats are endemic from the Hubei province in China and we discuss its potential role as reservoir for the evolution of SARS1 and SASR2.

## INTRODUCTION

The ongoing pandemic of COVID-19 (Coronavirus Disease-2019) has already had a deep health, economic and societal impact worldwide[1]. COVID-19 is caused by a novel betacoronavirus, SARS-CoV-2. Other members of *Coronaviridae* family include the highly pathogenic SARS-CoV and MERS-CoV, responsible for widespread outbreaks in 2002 and 2012, respectively[2].

SARS-CoV-2 encodes a large (30 kb) single stranded, positive sense RNA genome that contains multiple open reading frames (ORFs). ORF1a and 1ab produce two large replicase polyproteins precursors (450 kDa for ORF1a, 750 kDa for ORF1ab) which upon proteolytic cleavage generates 16 non-structural proteins (NSP), 1 to 16. Other ORFs encode the 4 main structural proteins of SARS-CoV2: spike (S), membrane (M), envelope (E) and nucleocapsid (N) proteins, as well as accessory proteins. Processing of the polyprotein precursors relies on the two viral proteases, NSP3 and NSP5. As shown in Figure 1A, NSP3 or papain-like protease (PLpro) is responsible for the proteolytic cleavage of nsp 1-4. The protein NSP5, or 3C-like protease (3CLpro), is responsible for the processing of other cleavage sites that results in nsp 5-16. NSP4 is uniquely cleaved by NSP3 on the N-terminus and NSP5 on the C-terminus[3].

**Figure 1:**
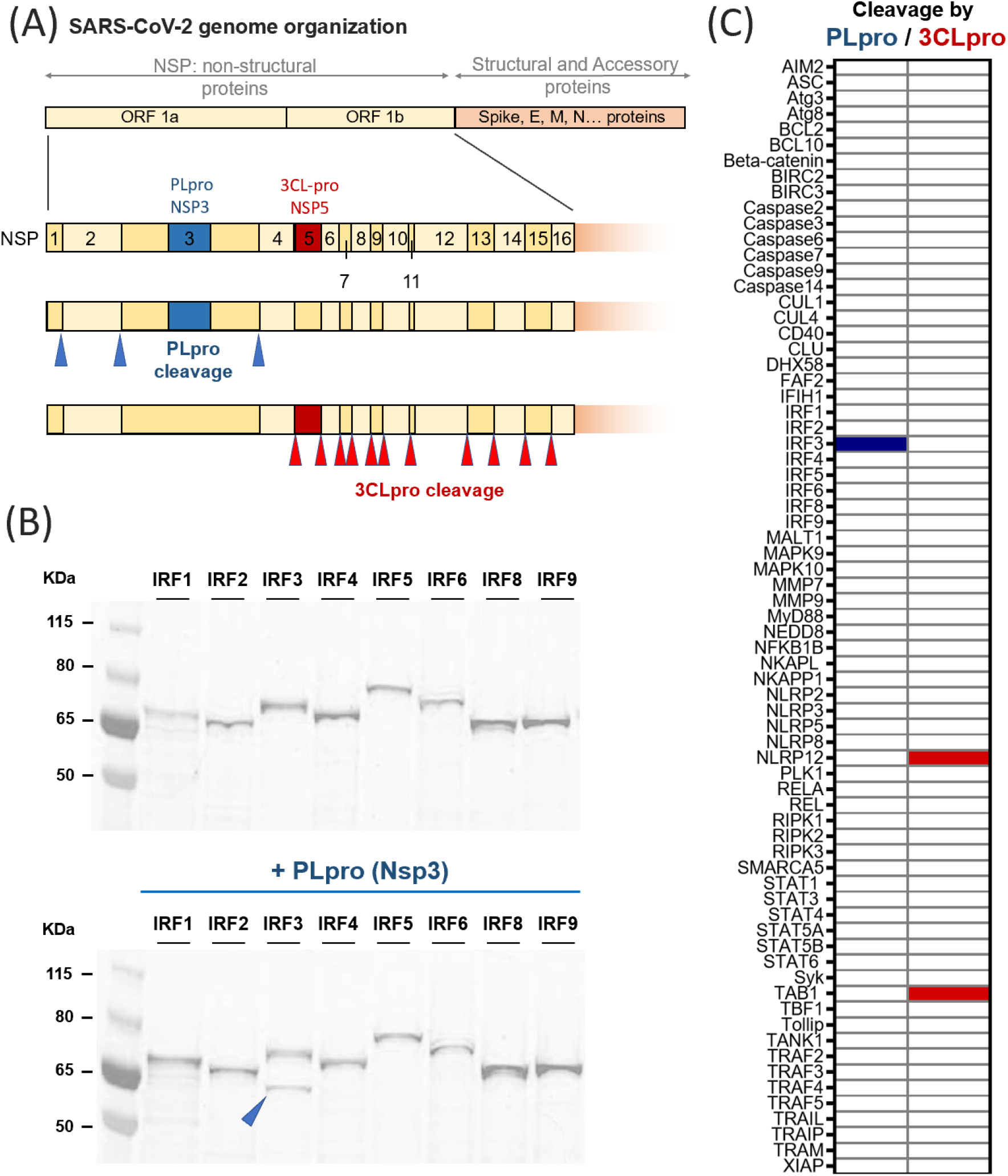
Principle of the screen of protease activity of SARS-CoV-2 PLpro and 3CLpro. (A) Schematic of the organization of the genome of SARS-CoV-2, focusing on the non-structural proteins Nsp1-16. As depicted, two proteases are encoded in ORF1a: NSP3 or papain-like protease (PLpro) and NSP5, or 3C-like protease (3CLpro). PLpro is responsible for three proteolytic cleavages, while the protein NSP5 cuts the large polyprotein at eleven different sites. (B) In our assay, the proteins are expressed using an eukaryotic cell-free system as a GFP fusion, and visualized using non-reducing SDS-PAGE using fluorescence. The proteases were added during protein expression, and the changes in banding pattern (protein size/integrity) were analysed to detect potential cleavage. Here, the results obtained for the family of IRF proteins are shown with PLpro, and the additional band obtained for IRF3 is indicated by a blue arrow (bottom panel, in the presence of NSP3/PLpro). (C) Overview of the proteins tested in this study and the proteolytic events detected: out of the 71 proteins tested, PLpro cleaves only IRF3 (indicated in blue) and 3CLpro cleaves NLRP12 and TAB1 (as shown in red).

As these two proteases are essential for viral replication, they are evident drug targets and considerable effort was spent by the structural biology community early in the SARS-CoV-2 outbreak. Structures of the protease domains of NSP3 and NSP5 have already been reported, opening the door to the identification or development of inhibitors, using virtual or high-throughput screening (e.g. [4–8]).

Viral proteins, especially those from RNA viruses that have stricter constraints on their genome size, often perform multiple tasks. In addition to performing their intrinsic functions in the viral life cycle, many have evolved to interfere with innate immune responses or otherwise co-opt the host cell’s machinery to facilitate optimal viral replication [9, 10]. The role of coronaviruses proteases in mediating virulence has been shown before [9, 11]. For example, PLpro of both SARS-CoV [12, 13] and MERS-CoV[14], as well as other coronaviruses[15, 16], lead to inhibition of the type I interferon pathway. Inactivation of different components of the pathway, including RIG-I[12], STING[12] TRAF3/ TRAF6[17], TBK1[16] and IRF3[13, 14, 18, 19], has been documented. These effects are mediated by partly mediated by the protease activity but mainly derive from the deubiquitinating and deIGSgylating functions associated with full-length NSP3[20–22]. SARS-CoV PLpro has also been reported to also activate TGF-β1 signalling[23] or down-regulate p53[24]. Similarly, 3CLpro from the feline coronavirus, feline infectious peritonitis virus (FIPV), inhibits type I interferon signalling through cleavage of NEMO[25], while the porcine deltacoronavirus (PDCoV) 3CLpro cleaves STAT2. SARS-CoV 3CLpro is responsible for virus-induced apoptosis[26]. Therefore, we hypothesized that these proteases could cleave human innate immune pathway proteins (HIIPs), leading to interference with or dysregulation of the host response.

To screen for HIIPs that might be targeted by SARS-CoV 2 PLpro or 3CLpro, we first leveraged the systems virology and systems biology tools present in relevant databases like InnateDB[27], PathBank [28] ViPR [29], VirHostNet2.0, and VirusMentha [30]to downselect a core set of HIIPs that covers almost all pathways involved in human innate immune responses. We then designed a fluorescent based *in-vitro* protease activity assay to screen this library of 71 full-length HIIPs (Fig. 1B). The protease domains of PLpro and 3CLpro of SARS-CoV2 were recombinantly purified and added to GFP-labelled target HIIPs, expressed in a eukaryotic cell-free system. Proteolytic cleavage was assessed by SDS-Page gels.

Our screen of 71 HIIPs (Fig. 1C) revealed that only 3 proteins were directly cleaved by these two viral proteases. Notably, we discovered that NSP3 directly cleaved IRF3, while NSP5 cleaved NLRP12 and TAB1. Surprisingly, both NLRP12 and TAB1 are cleaved at two different sites, creating three protein fragments. We identified the five cognate cleavage sites in these 3 HIIPs targeted by the PLpro and 3CLpro domains of SARS2 NSP3 and NSP5, respectively. Structure-function correlative analysis followed by comparative alignment of IRF3 and NLRP12 homologs across relevant mammalian orders reveal the potential explanatory power of our findings. The cleavage of IRF3 could explain the enigmatically blunted type-I IFN response that have been noted in early studies of SARS-CoV-2 infections[31, 32], while the NSP5 mediated cleavage of NLRP12 might explain the hyperinflammatory response linked to severe COVID-19 disease[33, 34]. Indeed, the lack or presence of cognate cleavage motifs in IRF3 and NLRP12 homologs presents interesting correlations with the presentation of disease in animal models; our results will enable the development of more effective animal models for severe COVID-19. Finally, we searched the available genomes of potential hosts, to determine whether SARS2 could have evolved into an animal where the different cleavage sites would be present. We found that out of 11 species of bats, only one presents all five cleavage sites identical to humans for NLRP12, TAB1 and IRF3. As *Myotis Davidii*, is found endemically in Hubei province of China, near the first epicentre of SARS-CoV-2 pandemic, we will discuss its potential role as reservoir host.

## RESULTS

### An *in-vitro* protease assay identifies targets of SARS-CoV2 PLpro and 3CLpro

The human target proteins to screen in the *in-vitro* assay were selected to contain major proteins associated with the signaling pathways of innate immunity and cell death[35]. Relevant to infection by viruses, proteins downstream of the nucleic acid sensors MDA-5 and RIG-I have been selected (e.g. MAVS, TRAF3, NFκB and IRFs). Also included are the effectors of the Toll-like receptors, TLR3 and TLR7, such as TRIF, TRAM, TRAF6 or TAB1. Proteins involved in cell-death (e.g. TRAF2, caspases, Bcl2, XIAP) were also included.

71 human proteins were cloned for expression as GFP-fusions in a cell-free expression system based on the eukaryotic organism of *Leishmania tarentolae* (LTE). This system produces full-length proteins up to 200 kDa with minimal truncations, minimal protein aggregation [36] and was previously used by our group to study the behaviour of various apoptotic proteins such as MyD88[37], MAL[38] or ASC and NLRP3[39].

The assay was designed as a one-pot reaction to rapidly identify proteolytic cleavage. Purified recombinant protease domains (adjusted to a final concentration of 10 μM) were added to the LTE during expression of the target proteins (see Supplementary Figure 1). The screening conditions were optimised to avoid off-target effects and false positives. The human proteins targets were typically expressed at low concentration (reaching at most 1 μM), in a crowded environment (LTE) that recapitulates the host cytosol. The proteases were allowed to react to target protein *de novo* synthesis for about 2 ½ hours, at 27°C (optimal temperature for protein expression using LTE). Under these conditions, it is probable that the activity of the proteases was greatly reduced.

We used the GFP-tag on the target protein to directly visualize cleavage using reducing SDS-PAGE (Fig. 1C). As expected, partial denaturation (i.e. no thermal denaturation) maintained the fluorescence of GFP so that proteins could be imaged without any subsequent purification steps. Comparing the protein migration patterns in the presence and absence of the protease identifies cleavable proteins. Indeed, an intact protein would appear on the gel as a single fluorescent band. If the protein is cleaved, then the gel will show either a single band, at a lower molecular weight (in the event of a complete proteolysis of all target proteins), or multiple fluorescent bands, corresponding to the full-length protein and its cleavage product in the case of an incomplete cleavage process, as described in Figure 1B. The use of a fluorescent tag also allows simple quantification of protein concentration based on fluorescence intensity[37, 39].

SDS-PAGE showed no difference of sizes in the presence or absence of viral proteases for most proteins, indicating that a large majority of human proteins were unaffected by the addition of PLpro and 3CLpro. This suggests that in our assay conditions, non-specific cleavage was not observed. However, 3 cases of proteolytic degradation were identified, giving confidence that the viral proteases are active. The fact that only specific members of the same family of proteins (e.g. IRFs, Fig. 1C) were cleaved (IRF3 cleavage by PLpro) suggests high specificity and the recognition of specific sequences. There was no common reactivity between PLpro and 3CLpro reinforcing the idea that each viral protease did indeed recognize a specific consensus sequence.

The screen results also revealed that protein expression levels were unchanged upon addition of the proteases, suggesting that none of the components required for cell-free expression were cleaved during the experiments. As shown in Supplementary Figure 2, the cell-free lysate acts as a crowded environment made up of many different proteins. Analysis of the Coomassie stained gels shows that staining intensity and profile were similar, even at the highest PLpro concentration, indicating that there was no significant cleavage of components of the cell-free reagent.

### PLpro selectively cleaves IRF3

To validate that PLpro could cleave IRF3, we titrated different concentrations of the protease in the reaction. As shown in Figure 2A, a strong concentration-dependence was observed, as expected. When the same experiment was performed in the presence of 3CLpro, no proteolysis was detected, validating that the cleavage is indeed specific to PL-Pro (Suppl. Fig. 3).

**Figure 2:**
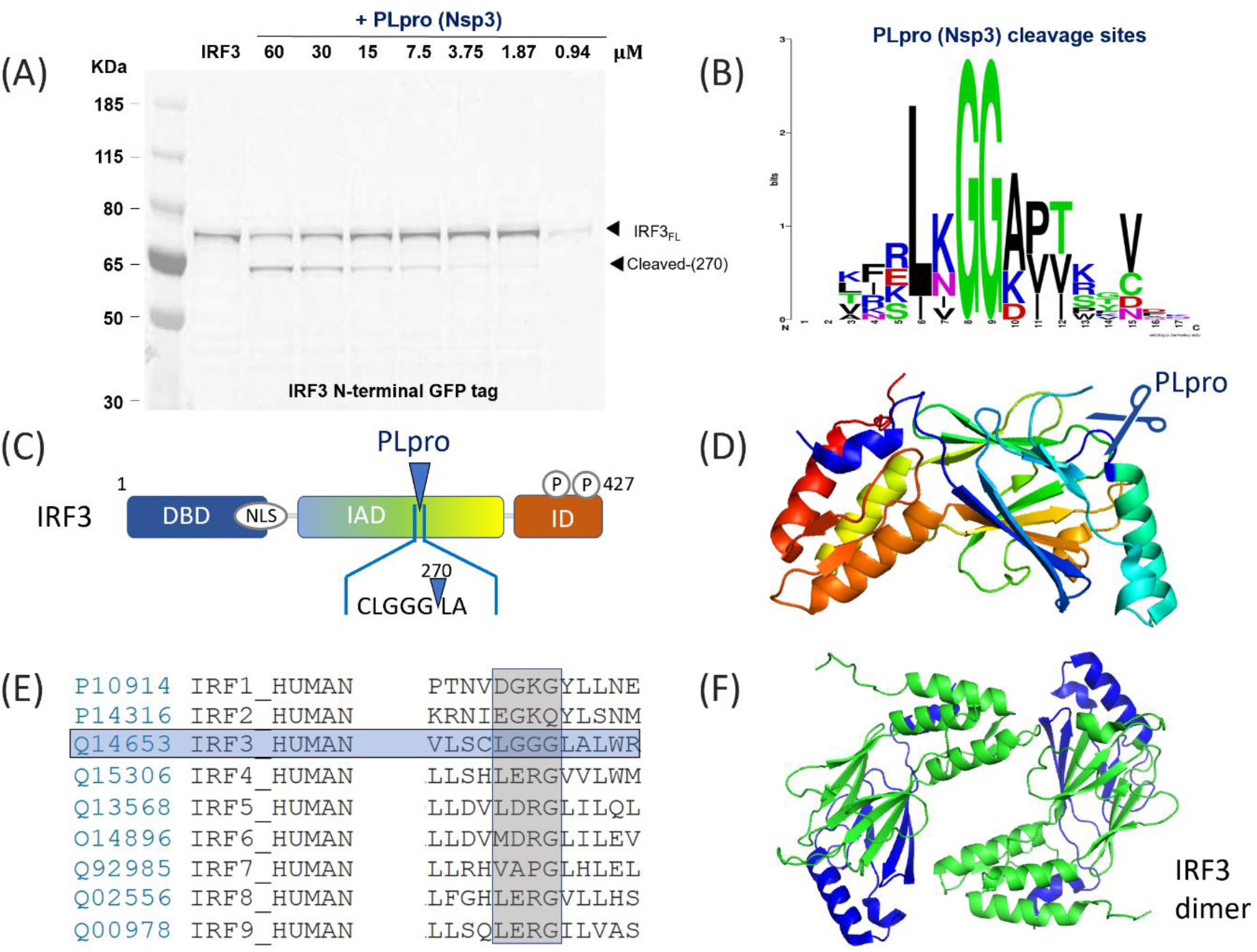
Cleavage of IRF3 by SARS-CoV-2 PLpro. (A): SDS-page analysis of the cleavage of human IRF3 protein, with a N-terminal GFP tag. The protein was expressed alone or in the presence of increasing concentrations of the SARS-CoV-2 protease PLpro. (B) Logo analysis of the cleavage site predicted from the polyprotein cleavage of SARS-CoV-2. Note that the first G is believed to be crucial for successful cleavage (C) representation of the domains found in IRF3 and the position of the cleavage site (D) Representation of IRF3 structure (from PDB 1J2F). The cleavage sequence LGGG is highlighted in blue; the site is presented in a flexible loop and seems fully exposed for cleavage by PLpro. (E) Alignment of the amino acids for human IRF1-IRF9, demonstrating that only IRF3 would be predicted to be cleaved as observed. (F) Structure of the IRF3 homodimer (from PDB 1QWT), showing the two fragments (green, N terminal, green, and blue for C-terminal). The cleavage seems to affect the dimeric interface, but this would need to be demonstrated experimentally.

We then set out to identify the cleavage site on IRF3. Based on the proteolysis sites on ORF1a and ORF1ab[3], and similarly to SARS-CoV PLpro [40] and MERS-CoV PLpro[41], SARS2-CoV PLpro recognises and cleaves after LxGG sequences (Fig. 2B). Sequence analysis of the human IRF3 shows the presence of a single LGGG sequence at residues 268-271 of the canonical isoform. Cleavage of the N-GFP tagged protein at this site would result in the formation of a GFP-tagged 57 kDa fragment (30 kDa +27 kDa for the GFP) and a 17 kDa untagged fragment, which corresponds well to the band obtained by SDS-PAGE (Fig. 2C). The identified cleavage site would be present on an exposed loop, based on previously solved structures (see Fig. 2D, PDB: 1J2F[42]) and therefore accessible to the protease. No such motif was found in any other member of the IRF family (Fig. 2E), in agreement with our data. On the contrary, recognition motifs present in other proteins of our test panel (LAGG in β-catenin, LVGG in STAT5A and LEGG in NLRP12) did not get processed by PLpro in our assay. In the case of STAT5A, the cleavage motif is partially buried in the protein (see Supplementary Figure 4). However, the LAGG motif in β-catenin is exposed at the surface in a structured a-helix (see Supplementary Figure 5); the LEGG motif in NLRP12 cannot be located on the only existing structure NLRP (NLRP3, PDB 6NPY). It is possible in addition to local structures, the residue between L and GG would contribute to the selectivity of PLpro cleavage.

IRF3 is a key mediator of type I interferon (IFN) response triggered by viral infections[43]. The C-terminal part of the protein is responsible for mediating interactions with upstream receptors and effectors STING, MAVS and TRIF[44]. IRF3 tail is also strongly targeted for post-translational modifications upon infection[45, 46], leading to its homodimerization (PDB: 1QWT, Figure 2F), translocation into the nucleus and transcriptional activation[47]. Therefore, we reasoned that PLpro cleavage of IRF3 would result in reduced IFN production, a feature that has been observed upon SARS-CoV2 infection[31].

### 3CLpro cleaves TAB1 and NLRP12

Similarly, we set out to validate 3CLpro proteolysis of TAB1 and NLRP12 (Fig. 3A and 4A). As before, we observed a concentration-dependent cleavage of both proteins and verified that PLpro did not have an effect, at any concentration (see Supplementary Figure 6 and 7).

**Figure 3:**
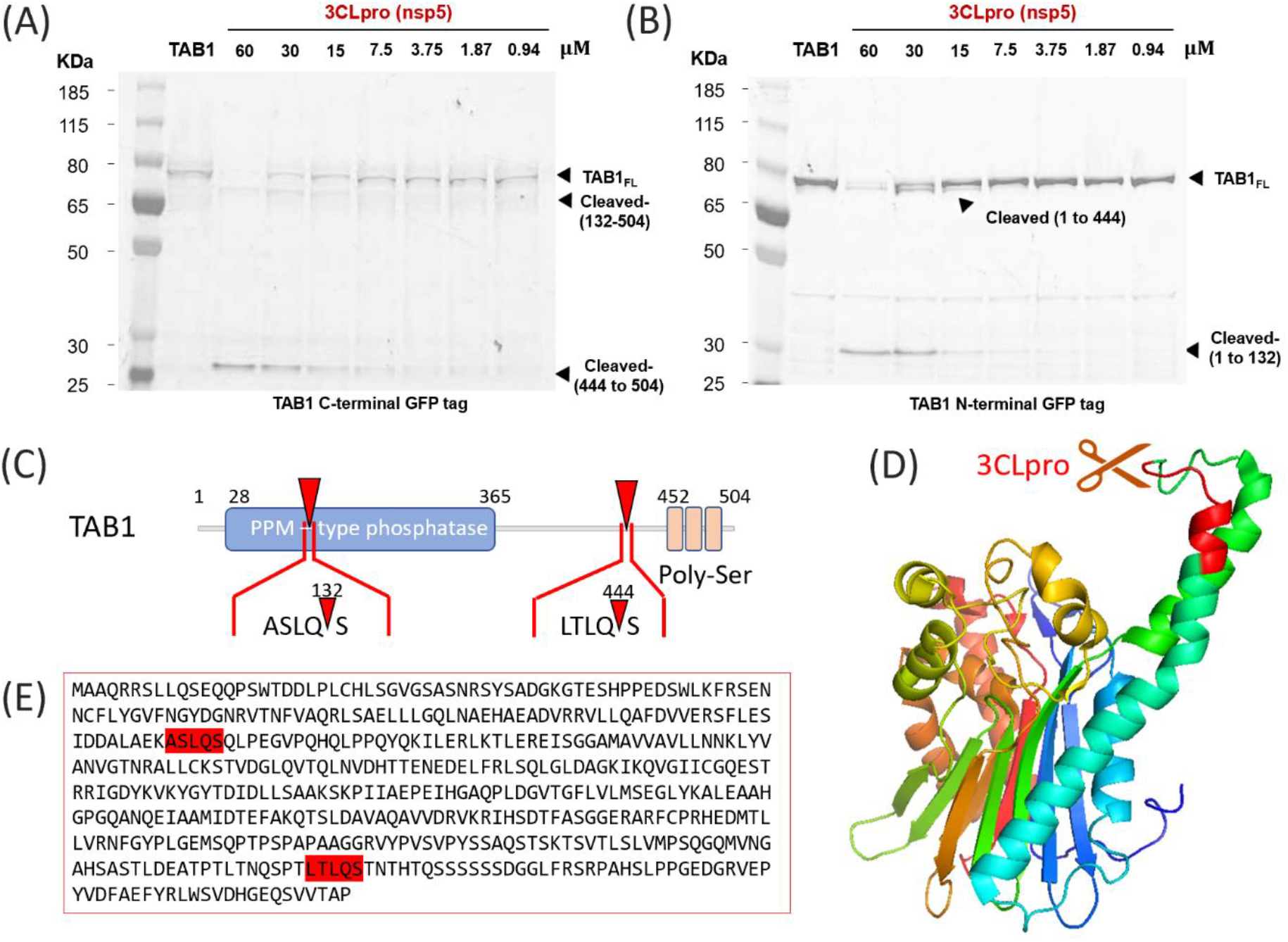
3CLpro (Nsp5) cleaves TAB1 at two separate sites. (A): SDS-page analysis of the cleavage of human TAB1 protein, with a C-terminal GFP tag. The protein was expressed alone or in the presence of increasing concentrations of the SARS-CoV-2 protease 3CLpro. The gel shows two additional bands upon cleavage, corresponding to the fragment 132-504 and to the fragment 444 to 504. These two fragments are visible as they carry the GFP tag; the fragments 1-132, 132-444 and 1-444 are not visible in this configuration. (B) same, but for the N-terminal GFP. In this case, the fragments 1-132 and 1-144 are fluorescent and can be detected on the gel, while the fragments 132-444, 444-504 and 132-504 are not fluorescent. (C) Schematic representation of TAB1 protein structure with the location of the identified cleavage sites on the primary sequence. (D) Representation of TAB1 structure (from PDB 2POM). The cleavage sequence ASLQS is highlighted in red; the site is presented in a flexible loop in an helix-loop-helix motif, and seems fully exposed for cleavage by 3CLpro. (E) full sequence of amino acids for human TAB1, showing the two putative cleavage sites ASLQS and LTLQS. The second cleavage site is not present in existing structures of TAB1 but is predicted to be flexible and accessible. Note that the removal of C-terminal tail of TAB1, is mimicked by a natural isoform.

**Figure 4:**
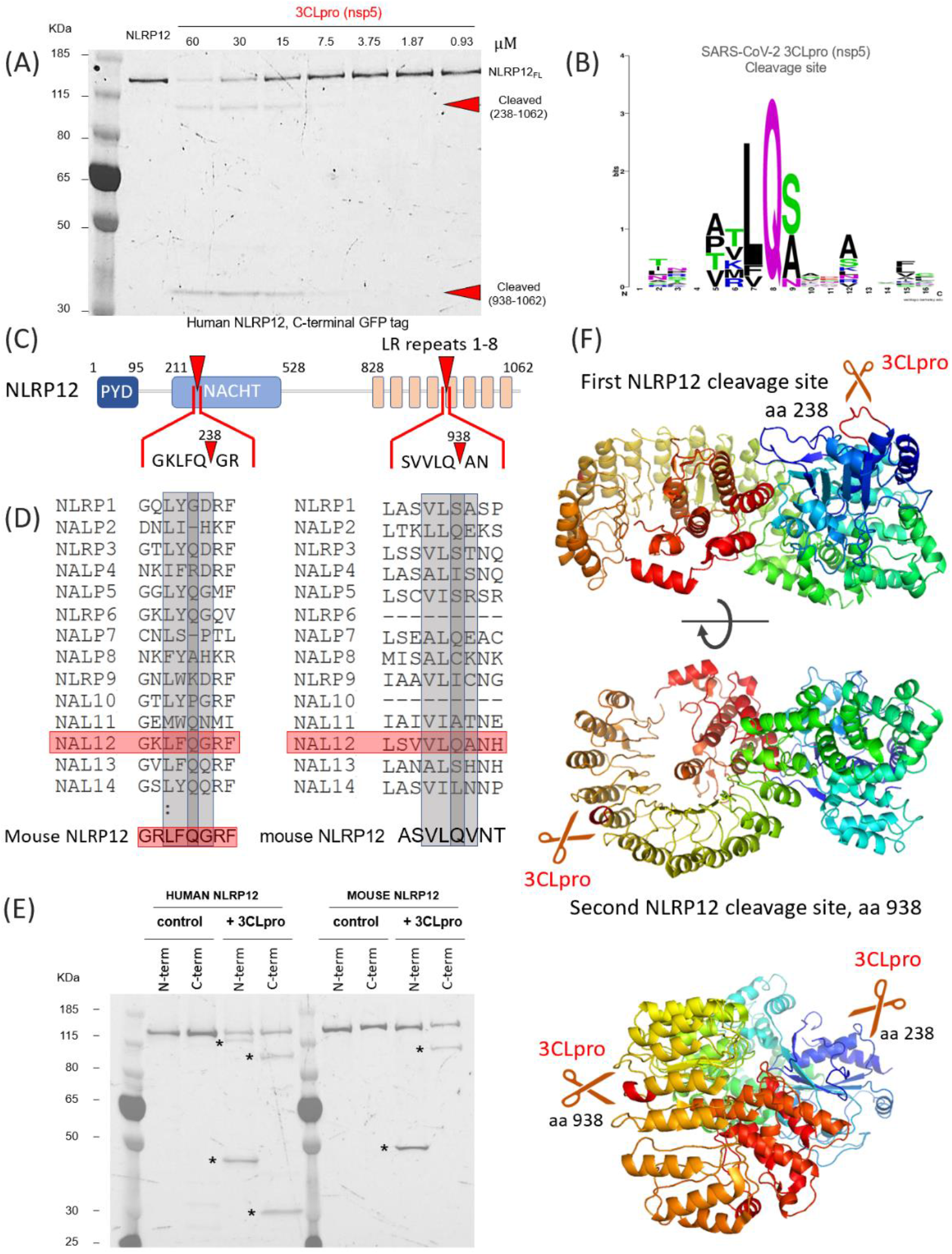
Cleavage of NLRP12 by 3CLpro of SARS-CoV-2. (A): SDS-page analysis of the cleavage of NLRP12 protein, with a C-terminal GFP tag. The protein was expressed alone or in the presence of increasing concentrations of the SARS-CoV-2 protease 3CLpro. The gel shows that two separate cleavage sites produce two distinct fragments attached to the C-terminal GFP (B) Logo analysis of the cleavage site predicted for 3CLpro, from the polyprotein cleavage of SARS-CoV- 2. Note that the Q residue in position P-1 is believed to be crucial for successful cleavage (C) Representation of the domains found in NLRP12 and the position of the cleavage sites.(D) Alignment of the amino acids for human NALPs (NLRPs), demonstrating that only NLRP12 would be predicted to be cleaved as observed. Below, alignment of mouse NLRP12, showing that the first cleavage site is conserved, but the second site presents an A→V mutation that would disrupt cleavage. (E) SDS-page analysis of human and mouse NLRP12, with different tag orientations, to demonstrate the differences between species. With an N-terminal GFP tag, the human NLRP12 appears as two fragments (1-238 and 1-938), as the fragments 238-938 and 938-1062 would be non-fluorescent. With a C-terminal tag, only the fragments 238-1062 and 938-1062 are fluorescent and are detected on the gel, and the fragments 1-238 and 238-938 are non- fluorescent. The banding patterns obtained in the presence of 3CLpro are consistent with the predicted sizes. Using the mouse NLRP12 constructs, only one cleaved fragment is observed in the N-term and C- term constructs. Here, a myc-mCherry was used and the tag was detected by the red fluorescence of mCherry. In the N-term configuration, a single cleaved product is detected, corresponding to the fragment 1-238. In the C-term configuration, the fragment 238-1062 is detected; this confirms that the LQA→LQV mutation found in mice inhibits cleavage by 3CLpro. (F) Representation of NLRP12 structure (derived from the structure of NLRP3 from PDB 6NPY). (top): The cleavage sequence LFQG (site 1, at residue 238) is highlighted in red; the site is presented in a flexible loop and seems fully exposed for cleavage by 3CLpro. (middle): The second cleavage site, (LQA) found at residue 938, it presented in a flexible loop and would be accessible at the tip of an alpha helix in the LR repeats. (bottom): in this view, both cleavage sites are visible.

SDS-PAGE analysis reveals the presence of two cleavage sites on TAB1, that can be more easily visualized when the GFP tag is placed at either the C-terminus (Fig. 3A) or the N-terminus (Fig.3B) of the protein. The recognition motif for 3CL-Pro of coronaviruses SARS-CoV[48], MERS-CoV[49] and SARS2- CoV[50] is often LQ/(S,A,G) although the protease can also cuts after FQ or VQ motifs[3]. Further, the presence of a combination of hydrophobic and positively charged residues at position P-3 and P-4 also seems to be preferred (Fig. 4B). In TAB1, we identified a LTLQS motif at position 441 of the canonical form, that would give rise to a 48 kDa N-terminal fragment and a 6 kDa C-terminal fragment. Another possible recognition motif (ASLQS) is present at position 129 and that would give a 14 kDa N-terminal fragment and a 40 kDa C-terminal fragment (Fig. 3C). Therefore, the two proteolytic fragments for C-GFP TAB1 correspond to amino-acids 133-504 and 445-504 (Fig. 3A) whereas N-GFP TAB1cleavage leads to the formation of proteolytic fragments corresponding to residues 1-132 and 1-444 (Fig. 3B). More details on calculation of sizes based on migration are included in Supplementary Figure 8 and 9. Based on the reported structure of TAB1 (Fig. 3D, PDB: 2J4O, [51]), the first cleavage site is on an exposed loop in the pseudo-phosphatase domain. The second cleavage site is in a disordered region of the protein that does not appear on the structure (see Figure 3E for the full sequence of human TAB1.)

Another target of 3CL-Pro revealed by our screen is NLRP12. NLRP12 is an intracellular pattern-recognition receptor, from the nucleotide-binding and oligomerization domain-like (NLR) receptor family, which regroups key mediators of the innate immune response and inflammation [52]. NLRP12 modulates the expression of inflammatory cytokines[53] through the regulation of the NFκB and MAPK pathways[54]. Formation of NLRP12 inflammasome can activate caspase 1[54, 55], which produces interleukin and leads to cell death. But NLRP12 can also form mixed inflammasomes with other NLRPs, including NLRP3, and then play an inhibitory role[56, 57]. NLRP12 is also involved in adaptative immunity and controls MHC class I expression through a yet ill-defined mechanism[58].

Unexpectedly, we also noted the presence of two additional bands when NLRP12 was submitted to 3CLpro treatment, suggesting that two cleavage sites would also be found in NLRP12 (Fig. 4A). In Figure 4B, we show the analysis of preferred residues for cleavage by 3CLpro of SARS-CoV-2. Sequence analysis identified a canonical LQA motif at residue 938, at the C-terminal of NLRP12, in the middle of the LR repeats (Fig. 4C). The cleavage at residue 938 would create a small C-terminal fragment (residues 938-1062), observed on the gel using the C-terminally GFP-tagged NLRP12 (Fig. 4A). There was no other canonical 3CLpro recognition sequence (LQS) in NLRP12, to explain the second proteolytic event. When expanding the search to degenerated sequences (FQ/VQ), we identified a KLFQG sequence at residue 238 (Fig. 4C). Cleavage at this site would result in the formation of a 93kDa C-terminal fragment (residues 241-1,062), observed in Figure 4A. Analysis of the sequences of other NLRPs (NLRP1-NLRP14) shows that both motives are unique to NLRP12 (Fig. 4D). Interestingly, the mouse homolog of NLRP12 possesses the first but not the second recognition motif (Fig. 4D). To further validate our data, we compared the effect of 3CLpro on human vs. mouse NLRP12, each tagged at the N- and C-terminal position. Figure 4E shows that human NRLP12 (left panel) is indeed cleaved twice, whereas mouse NLRP12 (right panel) is only processed once. Only the smaller N-terminal fragment and corresponding larger C-terminal fragment are conserved, indicating that cleavage at residue 238 occurs for both species, while cleavage at residue 938 is specific to human. The calibration of migration of the fragments on the SDS-page gels is further described in Supplementary Figure 10. Homology modelling of NLRP12, using the structure of its close relative NLRP3 (PDB: 6NPY[59]), shows that both sites are expected to be in an exposed, unstructured loops, readily accessible to a protease (Fig. 4F).

### Validation in SARS-CoV-2 infected cells

Similar to SARS, SARS2 utilizes ACE2 and TMPRSS2 to achieve its entry step [60–62]. The cleavage of IRF3, TAB1, and NLRP12 was demonstrated by *in vitro* PLpro or 3CLpro cleavage at the previous figures. To validate the cleavage effects upon SARS2 infection, we generated ACE2 expressing and ACE2/TMPRSS2 double expressing 293T cells by lentiviral expression system for enhancing the infectivity of SARS2. 293T-ACE2 and 293T-ACE2/TMPRSS2 cells were then infected with SARS2 and analyzed for viral and host protein levels. Fig. 5 shows that the expression of IRF3, NLRP12, and TAB1 (panels A-C, respectively) was decreased in virus-infected cells (lanes 2-3 and 5-6) compared to the mock-infected controls (lanes 1). The lower expression of ACE2 in 293T-ACE2/TMPRSS2 cells compared with that in 293T-ACE2 cells (lanes 4 and 1, respectively) might be caused by the proteolytic cleavage of ACE2 due to TMPRSS2 overexpression [63]; however, it did not affect the pattern of IRF3, TAB1, and NLRP12 during SARS2 infection. Combined with the results from *in vitro* assay, we found the proteases of SARS2 (PLpro and 3CLpro) can degrade IRF3, TAB1, and NLRP12 probably through their protease activity, leading to the imbalance responses of host innate immunity

**Figure 5:**
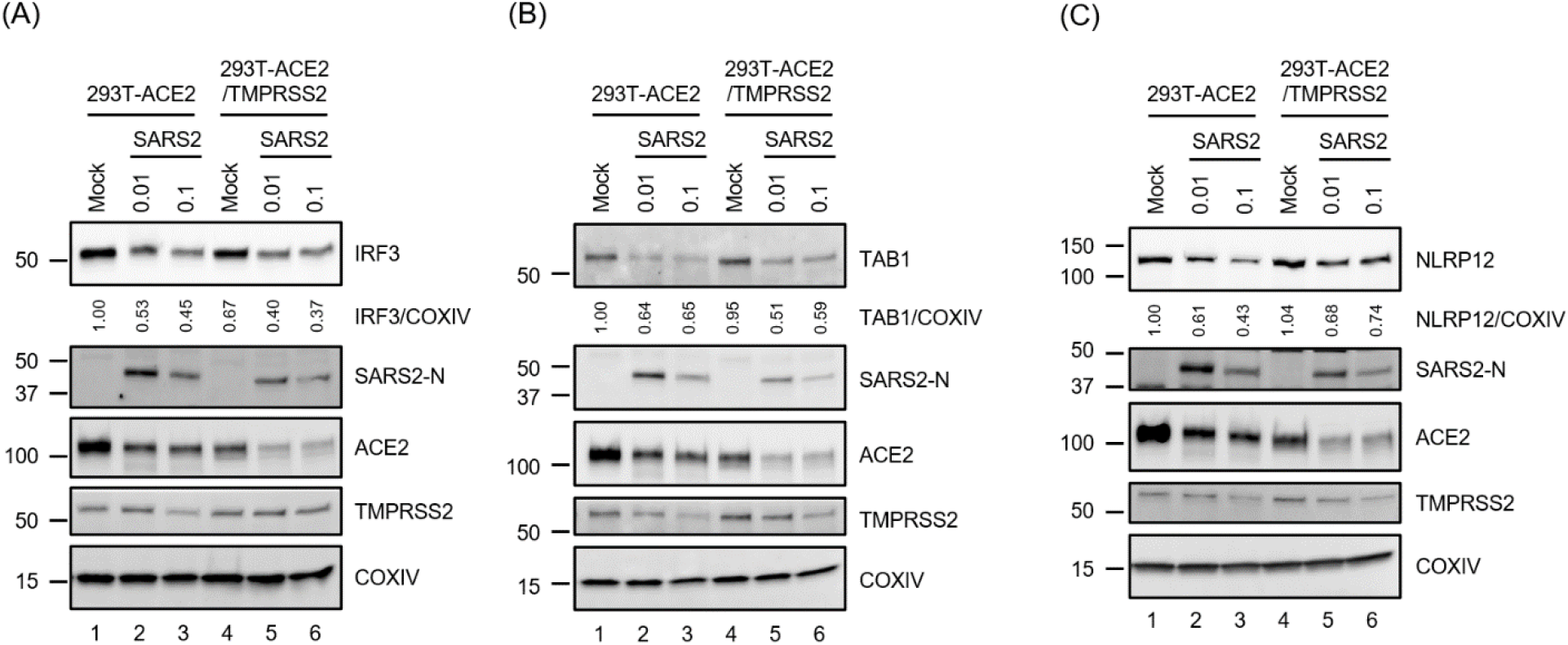
IRF3, TAB1 and NLRP12 is decreased upon infection with SARS-CoV-2 (SARS2) in 293T cells stably expressing human ACE2 alone (293T-ACE2) or with TMPRSS2 (293T- ACE2/TMPRSS2). (A-C) Stable 293T-ACE2 and 293T-ACE2/TMPRSS2 cells generated by lentiviral transduction were infected with SARS-CoV-2 at 0.01 or 0.1 MOI. Uninfected cells plated and treated identically served as mock-infected controls. 72 hpi, cell lysates were collected, and the indicated proteins were detected by western blot using the relevant antibodies as specified in methods. COXIV serves as the loading control. The relative amounts of IRF3, TAB1 and NLRP12 was quantified by densitometry and presented as a ratio of COXIV with the ratio obtained from mock infected 293T-ACE2 cells set at 1.

## DISCUSSION

### Experimental screens identify accessible cleavage sites in proteins and more importantly, detects non- canonical sequences

In this report, we show that the viral proteases PLpro and 3CLpro of SARS-CoV2 lead to the *in-vitro* proteolytic cleavage of three important proteins of the host immune response: IRF3, TAB1 and NLRP12 (Fig. 1B). These results first show the exquisite specificity of the viral proteases, with only 3 positive hits observed out of 142 experiments.

In the case of PLpro, the recognition sequence seems well defined and LxGG motifs are scarcely found in the proteins we studied. Nevertheless, of the 4 proteins that harbor such motifs, only 1 (IRF3) was cleaved *in vitro*. Indeed, the motif is present on an unstructured loop of IRF3 that made this site readily accessible. This shows that the presence of a recognition motif is required but not sufficient to predict biological activity. As such, experimental validation of potential targets, identified by bioinformatics remains critical. The identification of NLRP12 and TAB1 as substrates of 3CLpro further demonstrate the importance of this type of screening approaches. 3CLpro recognition motif is not as well defined as PLpro, and LQ/(S,A,G) motifs are ubiquitously found in the proteome. In our protein list, multiple *bona fide* recognition motifs based on LQ/S, LQ/A and LQ/G, respectively, can be found as well as several degenerated motifs. Yet only two targets (and four cleavage sites) were recognized by 3CLpro, emphasizing again the importance of site accessibility. More importantly, our data identify an unexpected cleavage site on NLRP12 (KLFQ/G) that does not resemble to the cleavage motifs present on the virus orf1a/1ab, further indicating that the determinants of selectivity for 3CLpro are yet to be identified.

### IRF3, TAB1 and NLRP12 are important components that drive the inflammatory response to SARS- CoV2 infection

Viral proteases have probably evolved to efficiently process their own polypeptides but their ability to target host proteins, especially the ones involved in host defence, provide them with an evolutionary advantage. The three HIIPs identified in this screen (IRF3, TAB1 and NLRP12) are important contributors to the innate immune and inflammatory responses, either driving or dampening it.

IRF3 belongs to the interferon-regulatory factor (IRF) family. All IRFs possess a well-conserved DNA-binding domain at the N-terminus and a variable C-terminal domain that mediates most of the interactions with the other IRF proteins and other co-factors[64]. Fine-tuning of the choice of partners rely on post-translation modifications, such as phosphorylation and ubiquitination, in this region. As mentioned before, IRFs, especially IRF1, 3, 5 and 7, are major contributors to the production of and response to type I interferons, which stimulate macrophages and NK cells to elicit anti-viral responses[47]. Therefore, IRF3 has been found to be targeted by several different classes of viruses. Paramyxoviruses, herpesviruses, reovirus and double stranded RNA viruses, to cite some examples, have been shown to interfere with IRF3 signalling through different strategies[65]. These include the control of protein expression, protein cellular localisation, modifications of the PTMs, inhibition of protein-protein interactions or induced cellular degradation[10]. Direct proteolytic cleavage of IRFs by viral proteases has also been identified before. Proteases of enteroviruses EV71[66] and EV-D68[67] have been shown to directly cleave IRF7.

Papain-like protease domains of several coronaviruses including SARS-CoV and MERS-CoV have been implicated in the observed reduction of IFN signalling upon viral infection[68]. Although this link seems well established, the molecular basis of this effect is still under debate. Whether the enzymatic activity is required is also controversial. Besides its proteolytic activity, PLpro of Coronaviridae also possess a deubiquitinase and deISGylation activity that contribute to inhibition of IRF3 activity[13, 18, 19, 69]. Further, SARS-CoV PLpro has been shown to target proteins upstream of IRF3. It can directly bind to STING, disrupt the formation of the STING-TRAF3-TBK1-IKK_ε_ complex or limit signalling by deubiquitinating STING, TRAF3, TRAF6 or TBK1[44]. Here we show that *in-vitro*, SARS-CoV2 PLpro was able to cleave IRF3 and validated that the effect could be observed in relevant virus infected cells (Figure 5). Direct proteolytic cleavage of IRF3 may contribute to the reduction of type I interferon production observed in COVID19 patients[31, 32].

TAB1 is part of the TAB1/2/3/TAK1 complex[70] that regulates the activity of TAK1 (TGFβ-activated kinase 1), in response to different stimuli including TGFβ, IL1, TNFα and upon viral and bacterial infection[71, 72]. TAK1 can then activate the NFκB pathway or signal through the MAP kinases pathway[73, 74]. Lei et al. showed that the 3C protease of enterovirus 71 (EV71) cleaved TAB1 (along with TAB2, TAB3 and TAK1) at two cleavage sites (Q^414^S and Q^451^S) to perturb the formation of the complex and inhibit cytokine release downstream of NFκB[66]. Our identified cleavage site, at Q^444^S, forms a protein that is reminiscent of the second isoform of TAB1 (TAB1 β, which lacks the C-terminus) that loses its interaction with TAK1 [75, 76]. Further, the poly-Ser region (452-457) is a substrate for p38 kinase[77] (whose binding site has been mapped to residues 408-414 on TAB1) and phosphorylation controls cellular localisation and activity of the protein[70]. Therefore, the loss of TAB1 C-terminus through 3CLpro cleavage would profoundly impact its ability to activate TAK1 and result in decrease production of cytokine through NFκB signalling.

Finally, we identified NLRP12 as a substrate of 3CLpro, and two cleavage sites could be identified. NLRP12, like other members of the NLRP family, possess a PYD domain, that binds the effector ASC through homotypic interaction, followed by a NACHT domain, which binds ATP and mediates activation of the protein, and a series of LR repeats that gives specificity to each member by modulating protein-protein interactions[78]. NLRP12, by similarity with NLRP3, is thought to be normally maintained in a monomeric, auto-inhibited conformation and release of this auto-inhibition is mediated by binding of a ligand (e.g. ATP) to the NACHT domain[79]. ATP binding to NLRP12 plays a major role in regulating the protein’s activity as it has been shown to not only induce self-oligomerisation, but also promote interaction with NFκB-induced kinase (NIK) and subsequent degradation of NIK[80]. Mutations in the ATP binding site are sufficient to increase production of proinflammatory cytokines and chemokines, mimicking loss of NLRP12[80]. The first cleavage site in NLRP12, at residue 238, is located the NACHT domain, in between the two walker motifs that mediate ATP binding (walker A; residues 217-224 and walker B: residues 288-299, by analogy with NLRP3[81]). This cleavage breaks the nucleotide binding site and whether this leads to activation or repression of the protein activity is an open question. Of note, this proteolytic cleavage also releases the PYD. PYD of ASC, the NLRP3/12 adaptor, has been shown to drive polymerization of ASC and formation of ASC specks upon inflammasome formation[82]. Whether NRLP12 PYD is able to polymerise on its own and induce downstream signalling and pyroptosis remains to be explored. The second cleavage site, at residue 938, releases 3 LRR motifs, and probably modifies protein-protein interactions. Most relevant to COVID-19, NLRP12 is known to negatively regulate the release of pro-inflammatory cytokines[83]. Mutations in NLRP12 [84] have been linked to autoinflammatory disorders, in particular the familial cold autoinflammatory syndrome 2 (FCAS2, OMIM: 611762). Of note, truncation R284X, close to the identified cleavage site has been identified in patient suffering from hereditary periodic fever syndrome and leads in vitro to a dramatic increase in NF-κB activation [85]. Up-regulation (and potentially over-activation) of NLRP12 has also been noted in patients with Kawasaki disease [86], a rare auto-inflammation of blood vessels in children. The emergence of Kawasaki-like syndromes in children positive for SARS-CoV 2 [87, 88] could point to a molecular link between NLRP12 cleavage and inflammatory sur-activation.

### Comparison between species can predict animal model accuracy to study particular effects of SARS-CoV2

The recognition sequences on IRF3 and NLRP12 are unique in their respective families of proteins, so we wanted to investigate how conserved these motifs were. To this end, we compared the protein sequences of IRF3 and NLRP12 among species, specifically around these sequences. In both cases, the cleavage sites were located on well conserved portions of the proteins but small differences at or near the cleavage site were identified. Figure 6 summarizes the conservation of the IRF3 and NLRP12 sequences for 21 species that could be use as infection models. The PLpro recognition sequence in IRF3 is present in most species, except for rodents and ferrets, where the G at P2 is replaced by K, R or N, which would not be permissive to proteolytic cleavage.

**Figure 6:**
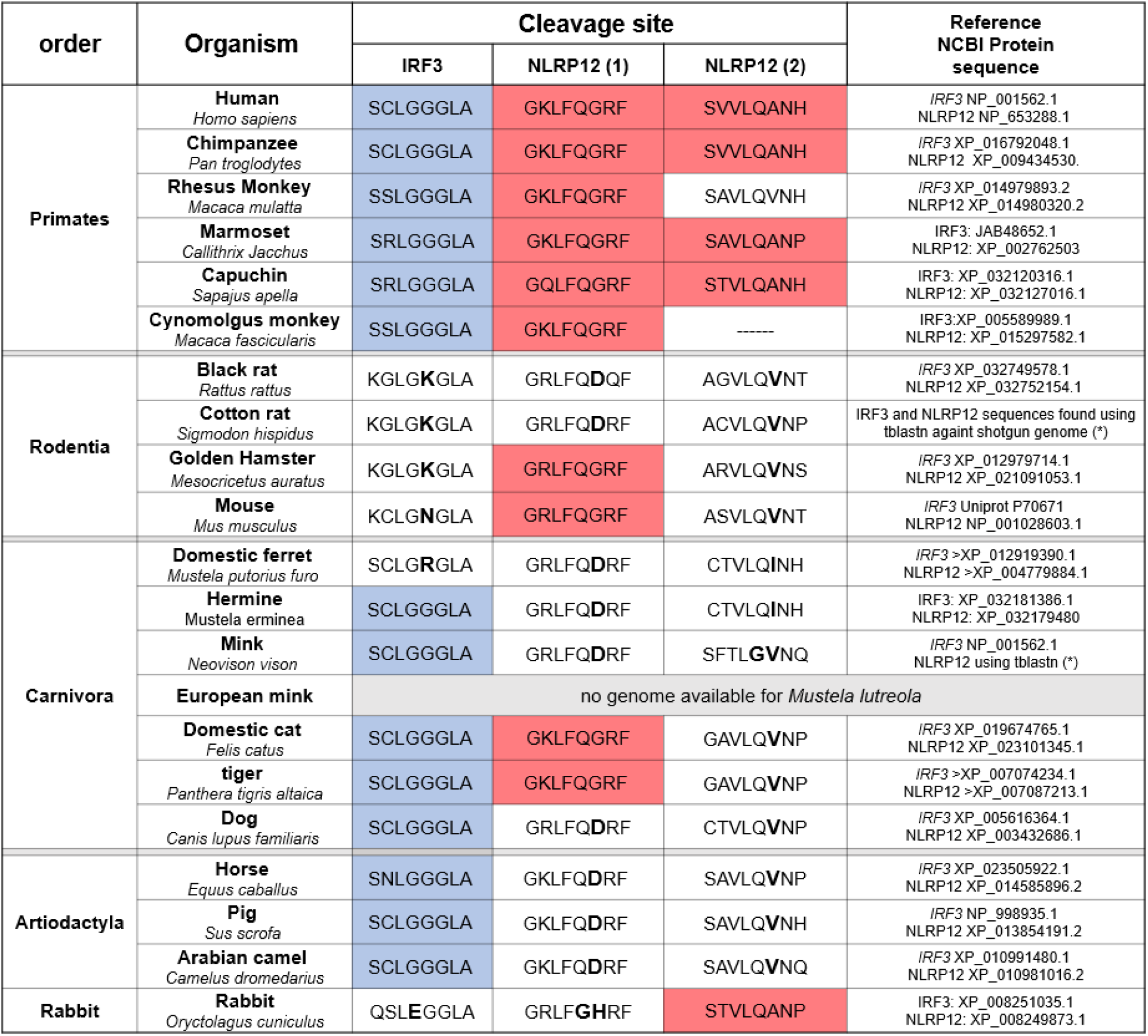
analysis of the protein sequences across species for IRF3 and NLRP12 cleavage sites. All primates tested presented both IRF3 and NLRP12 cleavage sequences, except for the Rhesus Monkey and Cynomolgus (“crab-eating”) monkey where the second NLRP12 cleavage site is mutated or missing. In rodents, most of the typical species used for testing in laboratories (rat, hamsters, and mouse) have a mutation in the most important residue for PLpro cleavage. Cats, tigers (and most feline, not shown) present both IRF3 and NLRP12 cleavage sites and we would predict that both proteins would be cleaved during SARS-CoV-2 infection. Ferrets have variations in these sites, and it is unlikely that the proteins would be affected. Cats and tigers can present respiratory symptoms, but dogs are not affected. Unfortunately, the genome of the European mink (*Mustela lutreola*) could not be found at this stage. European minks can present the disease and transmit SARS-CoV-2. Horses, pigs and camels possess the IRF3 cleavage site but not the NLRP12 sites; on contrary, the rabbit has a mutation in IRF3, but we would predict that the second NLRP12 site would be cleaved. Both cleavage sites of TAB1 are exactly conserved across all these species. (*) the protein sequences for Cotton rats (*Sigmodon hispidus*) and Minks (*Neovison vison*) were found using tblastn against shotgun genomes, with the Query AAH7172.1 for IRF3 protein [(Isoform 1) *Homo sapiens*] and NP_653288.1 for NACHT, LRR and PYD domains-containing protein 12 [(isoform 2) *Homo sapiens*].

We then examined the sequences of NLRP12. The motif around the first cleavage motif is mainly conserved, but we noticed that the amino acid directly after the cleavage site varied significantly. In primates, this amino acid is a small neutral amino acid (G) that is replaced by a bulkier, charged residue D or H in other species. A similar trend is observed for the second cleavage site where the small amino acid A found in human is replaced by larger amino acids (V or I). Such substitutions would likely affect the electrostatic environment and most likely inhibit the formation of active enzyme-substrate complexes. Interestingly, apart from primates, only cats and tigers have 2/3 similar recognition sequences to humans, for both proteins. It has been reported that, besides primates, cats are amongst the few species can could not only be infected with SARS-CoV2, but develop COVID-19 symptoms[89]. Anecdotal evidence suggest that Amur tigers (*Panthera tigris altaica*) [90] and European minks (*Mustela lutreola*) [91] could be infect/be infected by humans and develop symptoms. Unfortunately, the genome annotation for European minks is incomplete and does not allow for a comparison of its NLRP12 sequence.

This comparison once again highlights the difficulty of finding an animal model suitable for the study of SARS-CoV 2 infectivity and disease [92]. It has been rapidly established that mouse models of COVID-19 were ill-adapted as murine ACE2 the receptor for SARS-CoV2, is significantly different to the human isoform[93]. This led to efforts in developing transgenic mice models with humanized ACE2 which was successfully used to study infection by SARS-CoV [94] and SARS-CoV 2[95]. According to our analysis however, this model may not fully recapitulate the lack of interferon production if this is driven by IRF3 cleavage. We have also shown that mouse NLRP12 is only cleaved once, compared to two cleavage sites for human NLRP12 (Fig. 4D and 4E), although the functional impact of these cleavages remains to be studied. Ferrets are also a preferred model of respiratory viral infections and have been shown to be infected by SARS-CoV 2, but develop only mild symptoms [96]. So far, the most promising disease models amongst primates are rhesus macaques (*Macaca mulatta*) that develop pneumonia [97], although results of infection in capuchins (*Sapaju appella*) are yet to be published. In non-primates, cats present the most symptoms with massive lung lesions [89].

### Analysis of PLpro and 3CLpro cleavage in bats and other potential host species

We then extended our comparison to include wild animals that could have been host reservoirs of the virus, hypothesizing that the viral proteases had evolved to target innate immune proteins of their host. To date, the exact reservoir and intermediate hosts of the virus remain to be found. It is highly probable that the virus originates from a bat coronavirus (e.g. [98–100]) and the Malayan pangolins (*Manis javanica*) might have acted as intermediate hosts [101, 102] To this end, we compared the IRF3 and NLRP12 sequences in different bats, the Malayan pangolin and the Chinese tree shrew (*Tupaia chinensis*), based on data availability (Figure 7). Unfortunately, data for the masked palm civet (*Paguma larvata*), the intermediate host for SARS-CoV, could not be easily retrieved. As before, the IRF3 sequence around the cleavage site was remarkably conserved amongst species. On the contrary, the sequences for NLRP12 are highly divergent, especially at the C-terminus. Interestingly, this (limited) analysis identified only one species (*Myotis davidii*) that possess 3 similar cleavage sites compared to human.

**Figure 7:**
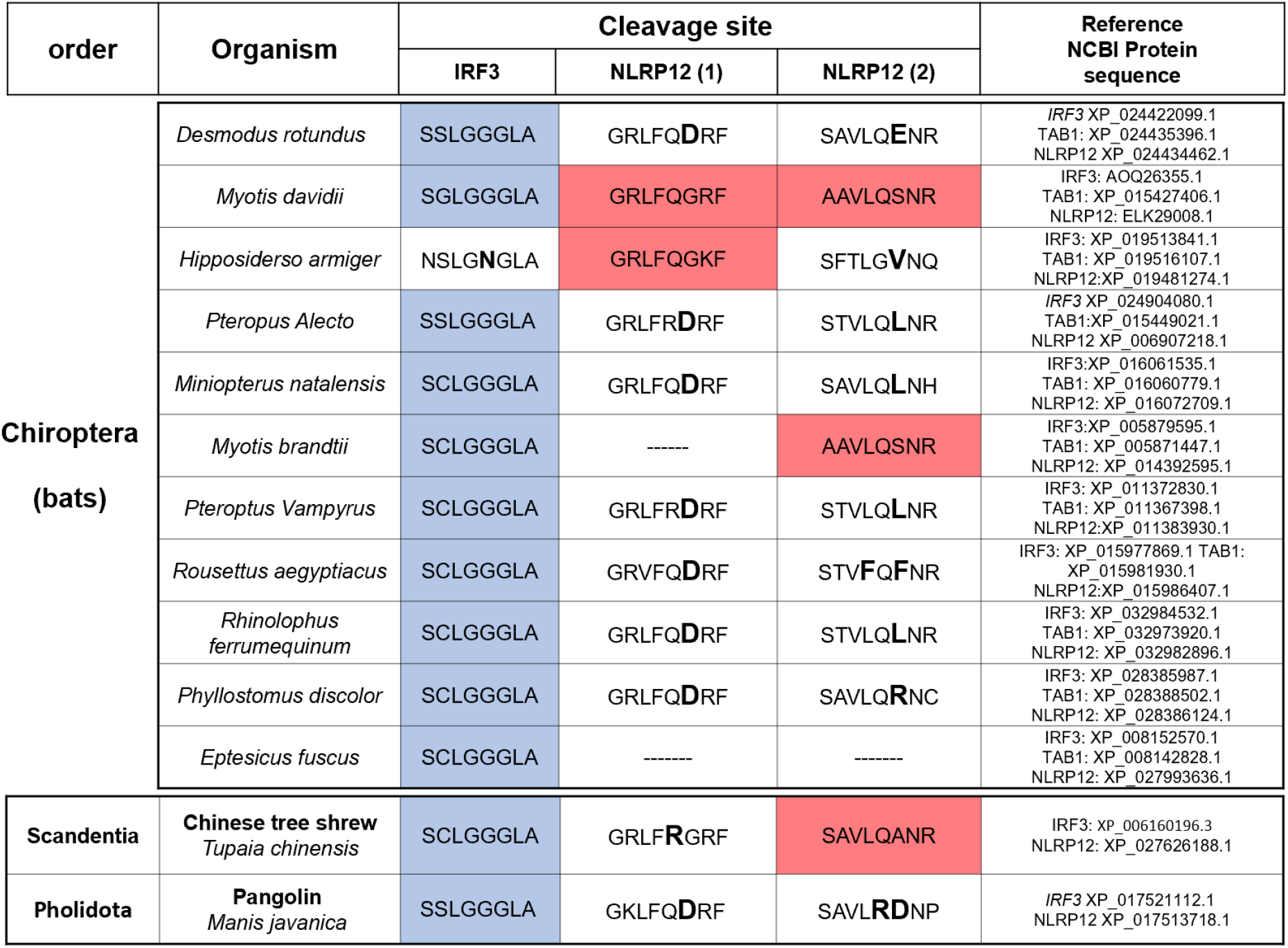
analysis of cleavage site sin potential host species. Most “exotic” species that would be relevant for SARS-CoV, MERS or SARS-CoV-2 present the correct cleavage site for IRF3. We found that one species of bats, Davids’ myotis, presents the three cleavage sites in IRF3 and NLRP12 (and also the two cleavage sites in TAB1). This species of bats is endemic to the province of Hubei, where the SARS-CoV-2 pandemic originated. Another small animal found in the same province, the Chinese tree shrew, displays at least two cleavage sites for IRF3 and NLRP12; the first cleavage site in NLRP12 (KLFRG) may potentially be cleaved. The species of pangolin described in China (Manis Javanica) does not possess the NLRP12 cleavage sites (note that surprisingly, an African pangolin presents all three cleavage sites identical to humans). Both cleavage sites of TAB1 are exactly conserved across all these species. The presence of the five human-like cleavage sites for IRF3, TAB1 and NLRP12 in a single species shows that it is possible that the SARS viruses could have gained the new functionality of cleaving these Human Innate Immune Proteins in a single reservoir host, potentially in *Myotis Davidii*.

## CONCLUSION

Overall, the method presented here enables medium to high-throughput screen of the activity of viral protein or bacterial effectors and will help design new antiviral and antibiotic strategies. In this study, we presented the results on the first 71 HIIPs (Human Innate Immune Proteins), and the screen will be expanded to cover more potential targets and pathways in the future. Our results show that in addition to the de-ubiquitinase activity of nsp3, SARS-Cov-2 uses its two proteases to further impact the host innate immune signalling. Our results were validated in SARS-CoV-2 infected A293T-Ace2 and 293T Ace2/TMPSSR2 cells, where decrease of IRF3, NLRP12 and TAB1 was demonstrated by Western Blotting. Our findings of IRF3 cleavage are consistent with the literature and the previous work showing an inhibition of interferon beta production in Covid-19 patients. More importantly, the direct cleavage of NLRP12 by 3CLpro could explain the hyper-inflammation observed in some patients.

The fact that the two proteases of SARS-CoV-2 could have evolved to interfere with innate immunity is attractive. If one considers that most of evolution is driven by lucky side-effects, the additional targets IRF3, TAB1 and NLRP12 could give a selective advantage if their cleavage would either enhance transmission or delay the response to infection. The discovery that all five cleavage sites are present in a single species of bats gives support to this theory. The fact that *Myotis davidii* can be found near the epicentre of the SARS2 pandemic makes it a possible candidate for a previous reservoir host, even if it does not exclude other hosts for SARS1 and SARS2. Further studies will be required to determine if these hosts display positive selection around the cleavage sites, and if zoonosis can be more precisely dated and determined.

## ACKNOWLEDGEMENTS

The authors would like to thank Katherina Michie, Jack Bennett and key personnel at the protein production facility of UNSW for the purification of PLpro and 3CLpro of SARS-CoV-2. The authors would like to thank Dr Kate Schroeder for the gift of the NLRP12 mouse plasmid in a previous collaboration, and Prof. Alexandrov for the cell-free plasmids compatible with LTE protein production.

MM and EO performed data collection for cell-free HIIPs cleavage experiments; H-PC performed SARS- CoV-2 infection experiments, PRS performed data collection of cell-free HIIPs, AB and DH contributed to the manufacture of the in-house LTE reagents and supplementation of the extracts, AF supervised collection of SARS-CoV-2 infection data and contributed to design of the experiments. DJ designed the constructs for NSP3 and NSP5, supervised their purification and wrote the manuscript. BL contributed to the design of the experiments, supervised the data collection on SARS-CoV-2 infected cells, and wrote the manuscript. ES and YG designed the project, supervised cell-free experiments on HIIPs and wrote the manuscript.

## MATERIALS AND METHODS

### Purification of PLpro (nsp3) and 3CLpro (nsp5)

E. coli C41(DE3) cells were transformed with the SARS-CoV-2 pET-28a-nsp5 plasmid (IDT DNA). Single colonies were used to inoculate LB media supplemented with kanamycin (50 μg/mL). 500 mL cultures were grown at 37°C until an OD600 of 0.6 was reached, cooled to 28°C and induced for 4 hours with 1 mM IPTG. Following growth, the cells were pelleted by centrifugation (5000 x g, 20 min, 4°C) and washed with 1X PBS before storage at −80°C. Frozen cell pellets were resuspended in buffer A (20 mM sodium phosphate buffer pH 8, 500 mM NaCl, 10 mM imidazole) and lysed on ice via sonication using a Branson SFX250 Sonifier (8 minutes at 50% amplitude, 2 s pulse on, 2 s pulse off). Cell debris was removed from the lysate by centrifugation (10,000 x g, 30 min, 4°C) and subsequent filtration of the supernatant through a 0.22 μm syringe filter. 20 mL of the clarified lysate was loaded onto a 1 mL HiTrap IMAC Sepharose FF column (GE Healthcare, Illinois) charged with Ni2+ and preequilibrated with buffer A. Unbound proteins were removed from the column through washing with 10 column volumes (CV) of buffer A. Bound proteins were eluted with a stepwise gradient of buffer B (20 mM sodium phosphate buffer pH 8, 500 mM NaCl, 500 mM imidazole) as follows: 0-5% B, 2 CV; 5% B, 5 CV hold; 5-25% B, 2 CV; 25%B, 5 CV hold; 25-100% B, 2 CV; 100% B, 5 CV hold. Fractions containing nsp5 were exchanged and concentrated into buffer C (20 mM Tris, pH 7.4, 10% glycerol), flash frozen and then stored at −80°C. All purification steps were performed on ice or at 4°C. Protein concentrations were determined using a linearized Bradford protein assay.

### Cloning and expression of the HIIPs

The 71 Human Innate Immune Proteins (HIIPs) listed in Figure 1C were cloned as GFP or mCherry fusions into dedicated Gateway vectors for cell-free expression. Open Reading Frames (ORFs) were sourced from the Human ORFeome collections, versions 1.1, 5.1 and 8.1 and transferred into Gateway destination vectors that include N-terminal or C-terminal Fluorescent proteins [103]. Most proteins screened were expressed as N-terminal enhanced GFP fusions (vector pCellFree G03); for TAB1 and NLRP12, C-terminal GFP were also used to validate the cleavage sites. The specific Gateway vectors were created by the laboratory of Pr. Alexandrov and sourced from Addgene (Addgene plasmid # 67137; http://n2t.net/addgene:67137; RRID:Addgene_67137). Mouse NLRP12 constructs were sourced from the laboratory of Dr Kate Schroeder (IMB, University of Queensland).

The HIIPs were expressed in vitro using a cell-free expression system derived from *Leishmania tarentolae* [104, 105]. This eukaryotic system enables expression of full-length proteins with minimal truncations and non-specific aggregation, for proteins up to 150kDa in size []. This system has been used recently to study the folding and oligomerisation of NLRP3 proteins and the polymerization of ASC [REF JMB] or the formation of higher-order assemblies of MyD88 [REF bostjan REF BMC]. The expression is simply set-up as a one-pot reaction where the plasmid encoding the protein of interest is added to the *Leishmania tarentolae extracts* (LTE); expression occurs within 3h at 27°C and expression yields can be evaluated by the fluorescence intensity of the GFP/mCherry tags [106].

### Detecting proteolytic cleavage of the HIIPS

The 71 HIIPs proteins were expressed individually in 10 μL reactions (1μL DNA plasmid at concentrations ranging from 400ng/μL to 2000ng/μL added to 9 μL of LTE reagent). The mixture was incubated for 30 minutes at 27°C to allow the efficient conversion of DNA into RNA. The samples were then split into controls and protease-containing reactions. The proteases PLpro (nsp3) and 3CLpro (nsp5) were added at various concentrations, and the reactions were allowed to proceed for another 2.5h at 27°C before analysis.

The controls and protease-treated LTE reactions were then mixed with LDS (Bolt LDS Sample Buffer, ThermoFisher) and loaded onto SDS-page gels (4-12% Bis-Tris Plus gels, ThermoFisher); the proteins were detected by scanning the gel for green (GFP) or red (mCherry) fluorescence using a ChemiDoc MP system (BioRad) and proteolytic cleavage was assessed from the changes in banding patterns, as shown in Figure 1B. Note that in this protocol, the proteins are not treated at high temperature with the LDS and not fully denatured, to avoid destruction of the GFP/mCherry fluorescence. As proteins would retain some folding, the apparent migration on the SDS-page gels may differ slightly from the expected migration calculated from their molecular weight. We have calibrated our SDS-page gels and ladders using a range of proteins, as shown in Supplementary Information.

### Preparation of LTE system

*Leishmania tarentolae* extracts were prepared in house using the protocol described previously [104, 107] check REF]. Briefly, *Leishmania tarentolae* Parrot strain was obtained as LEXSY host P10 from Jena Bioscience GmbH, Jena, Germany and cultured in TBGG medium containing 0.2% v/v Penicillin/Streptomycin (Life Technologies) and 0.05% w/v Hemin (MP Biomedical). Cells were harvested by centrifugation at 2500 x g, washed twice by resuspension in 45mM HEPES, pH 7.6, containing 250mM Sucrose, 100mM Potassium Acetate and 3mM Magnesium Acetate and resuspended to 0.25g cells/g suspension. Cells were placed in a cell disruption vessel (Parr Instruments, USA) and incubated under 7000 KPa nitrogen for 45 minutes, then lysed by rapid release of pressure. The lysate was clarified by sequential centrifugation at 10 000 x g and 30 000 x g and anti-splice leader DNA leader oligonucleotide was added to 10μM. The lysate was then desalted into 45mM HEPES, pH 7.6, containing, 100mM Potassium Acetate and 3mM Magnesium Acetate, supplemented with a coupled translation/transcription feeding solution and snap-frozen until required. We verified that the expression patterns and cleavage of the proteins in this study was independent of the batch of LTE used.

### Western blot and antibodies

Cells were lysed with RIPA lysis buffer (Pierce) containing a cocktail of protease inhibitors (Cell Signaling). Equivalent amounts of proteins determined by the Bradford protein assay (Bio-Rad) were separated by SDS-PAGE and transferred through the Trans-Blot Turbo transfer system (Bio-Rad). To avoid the nonspecific antibody reaction, the membranes were blocked with Intercept Blocking Buffer (LI-COR) prior to the addition of primary antibodies. After primary antibodies incubation, the blots were then treated with Alexa Fluor 647 conjugated secondary antibodies (Invitrogen) and developed signals using the ChemiDoc MP image system (Bio-Rad). The following commercial antibodies were used: anti-IRF3 (ab68481) and anti-ACE2 (ab108252) from Abcam, anti-NLRP12 (PA5-21027) from Invitrogen, anti-TAB1 (#3226) from Cell Signaling, anti-SARS-CoV-2 N (GTX632269) from Genetex, anti- TMPRSS2 (sc-515727) from Santa Cruz Biotechnology, anti-COXIV (11242-1-AP) from Proteintech.

## SUPPLEMENTARY INFORMATION

**Supplementary Figure 1:**
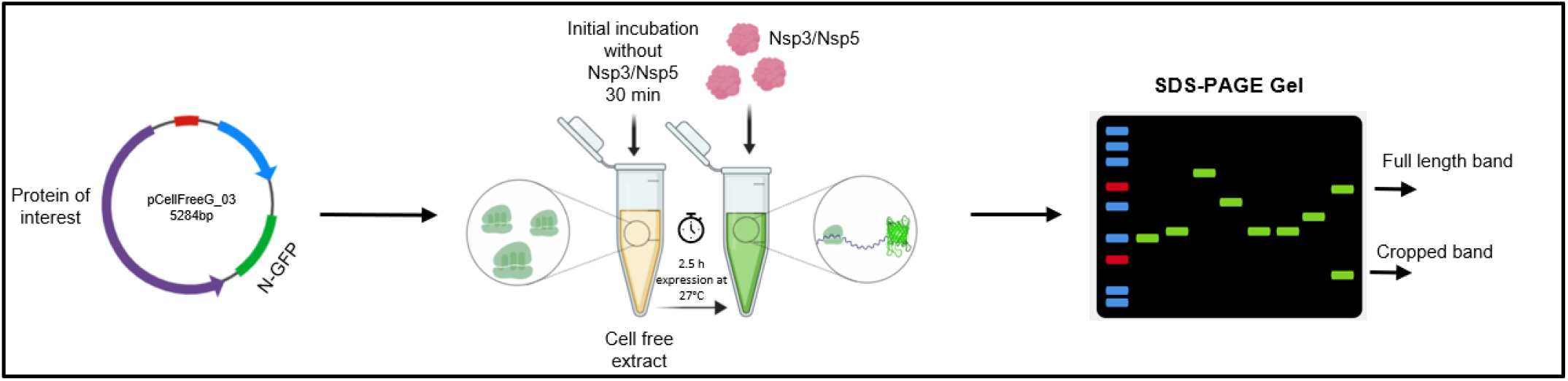
screen of proteolytic cleavage of Human Innate Immune Proteins (HIIPs) Open Reading Frames (ORFs) encoding 71 human innate immune proteins were cloned into Gateway vectors for cell-free expression as GFP-fusions. After mixing with the PLpro or 3CLpro proteases of SARS-CoV-2, the proteins were expressed for 2 ½ hours and analysed by SDS-page gels. The cleavage of the HIIPS creates additional fluorescent bands in the migration of the GFP-tagged proteins.

**Supplementary Figure 2:**
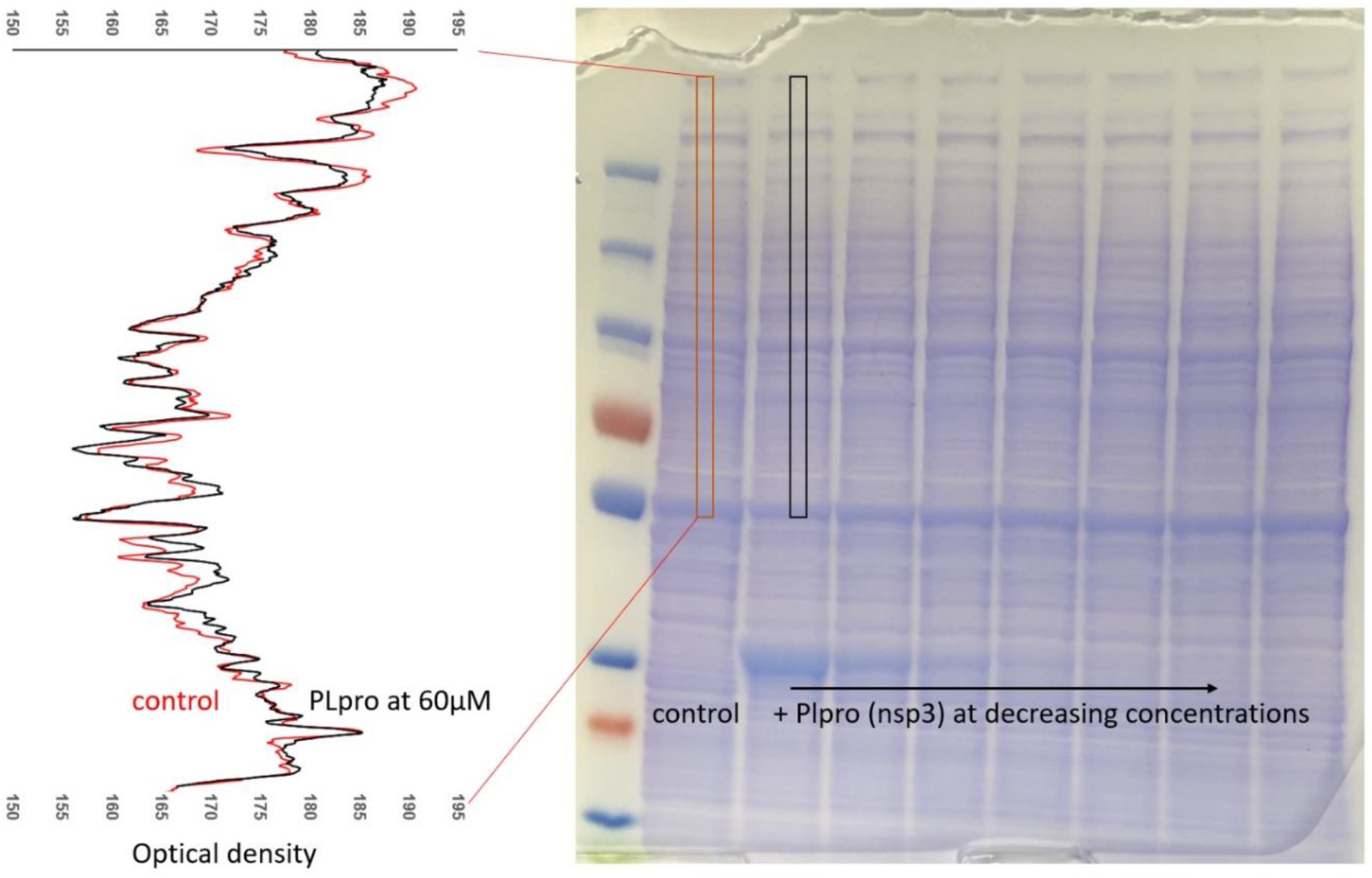
controlling for non-specific cleavage of proteins from LTE expression system. The cell-free expression system was loaded onto the SDS-page gels following the classic protocol and stained by Coomassie to reveal the proteins present in the LTE system. The SARS-CoV-2 protease PLpro (nsp3) was added at concentration ranging from 60 μM to 1μM (same as in Figure 2A). The gel shows on the right that the banding pattern of LTE is not affected by PLpro. (left): the density of bands across the control (LTE without PLpro added) was compared to the one obtained when 60μM of purified PLpro was added in the expression system. No significant differences were noted, confirming that PLpro doesn’t have a non-specific cleavage activity on the proteins of LTE. Also, the intensity of the GFP-tagged protein bands on the gels did not vary significantly when PLpro or 3CLpro were added, suggesting that the expression levels were unaffected, and that the components of the cell-free systems essential for expression were not cleaved.

**Supplementary Figure 3:**
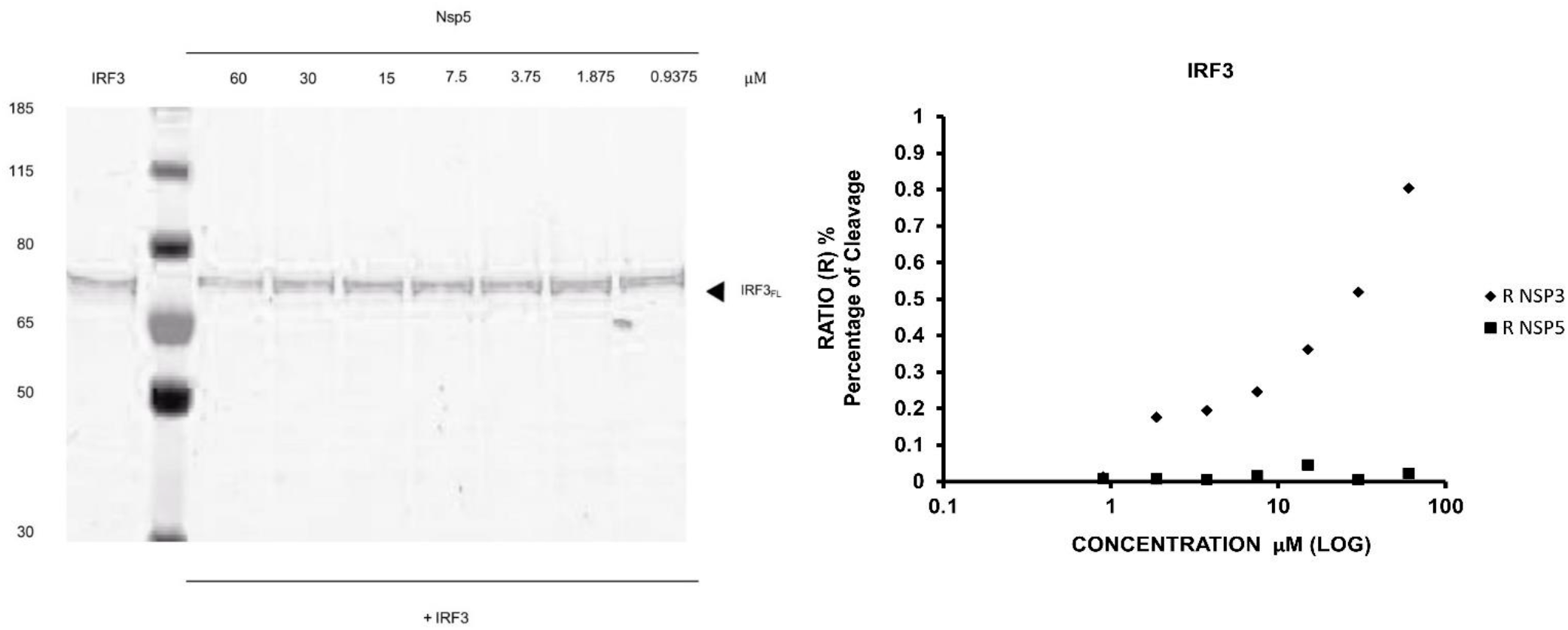
quantification of IRF3 cleavage by NSP3 and NSP5. (left): SDS-page analysis of the cleavage of IRF3 protein, with a N-terminal GFP tag. The protein was expressed alone or in the presence of increasing concentrations of the SARS-CoV-2 protease PLpro (nsp3). The gel shows no cleavage site. (right): Dose-response curve of SARS-CoV-2 protease PLpro (nsp3) and 3CLpro9 (nsp5) on Human IRF3. The result from the SDS-page gel was processed using IMAGE J and the Percentage of cleavage (Ratio R) was calculated by measuring the length of the line, in pixels.

**Supplementary Figure 4:**
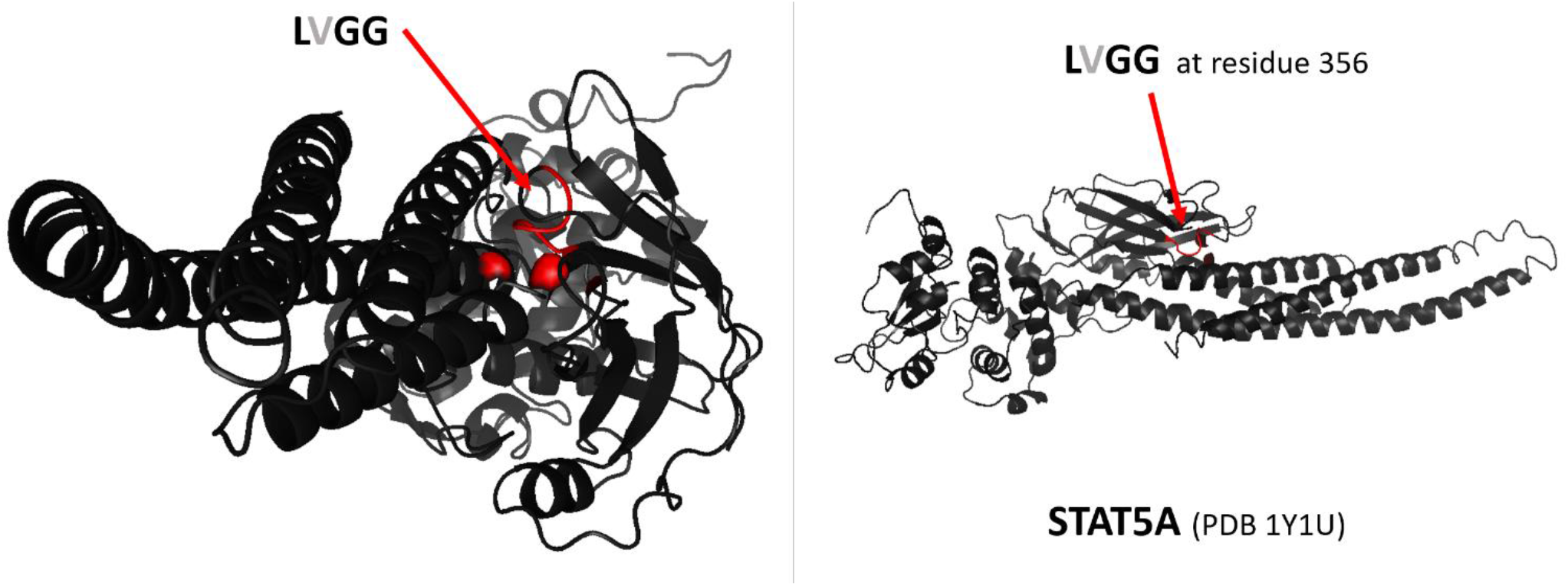
localization of a putative cleavage sequence in STAT5A. In our experiments, the protein STAT5A is not cleaved by PLpro or 3CLpro of SARS-CoV-2

**Supplementary Figure 5:**
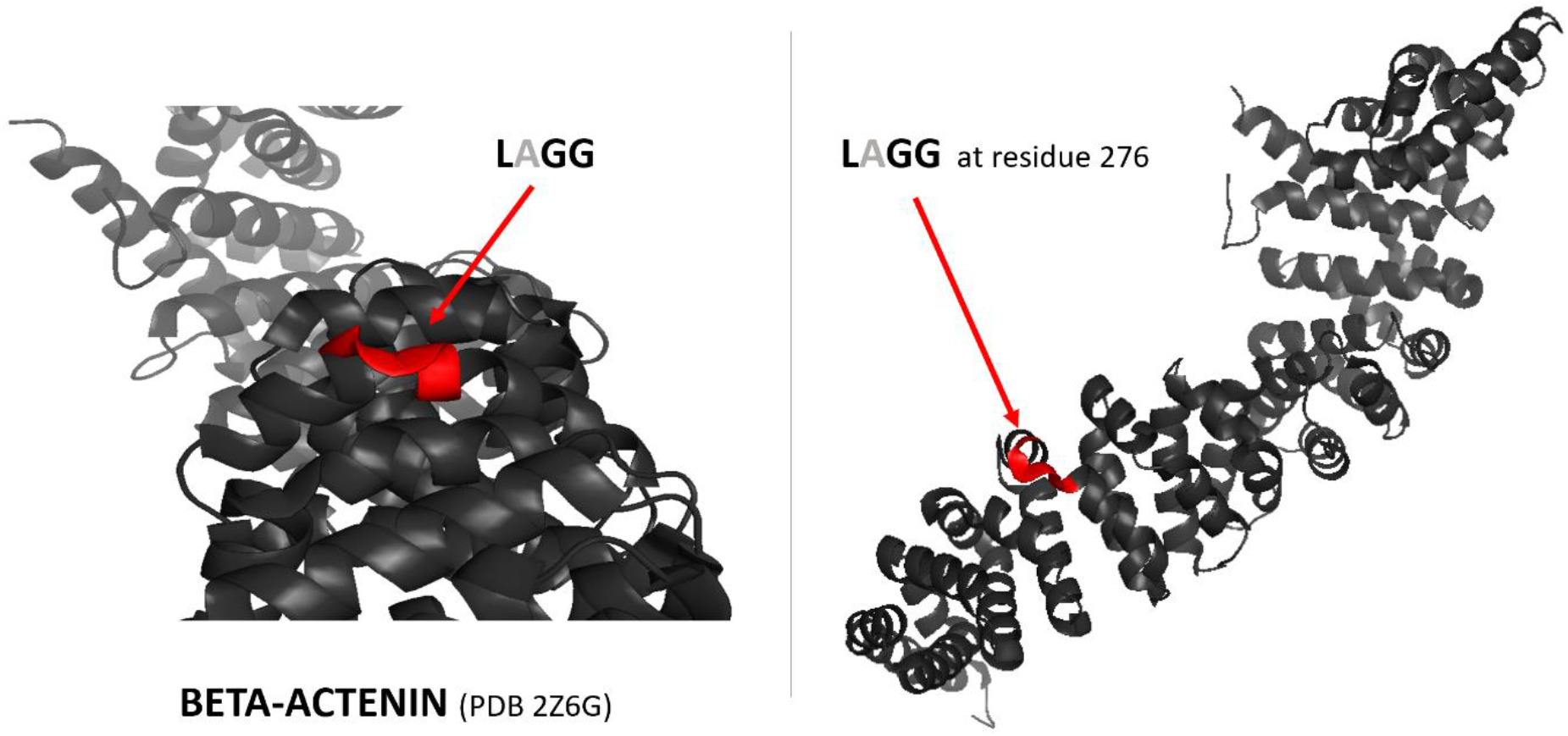
localization of a putative cleavage sequence in beta-Catenin. In our experiments, the protein beta-Catenin is not cleaved by PLpro or 3CLpro of SARS-CoV-2

**Supplementary Figure 6:**
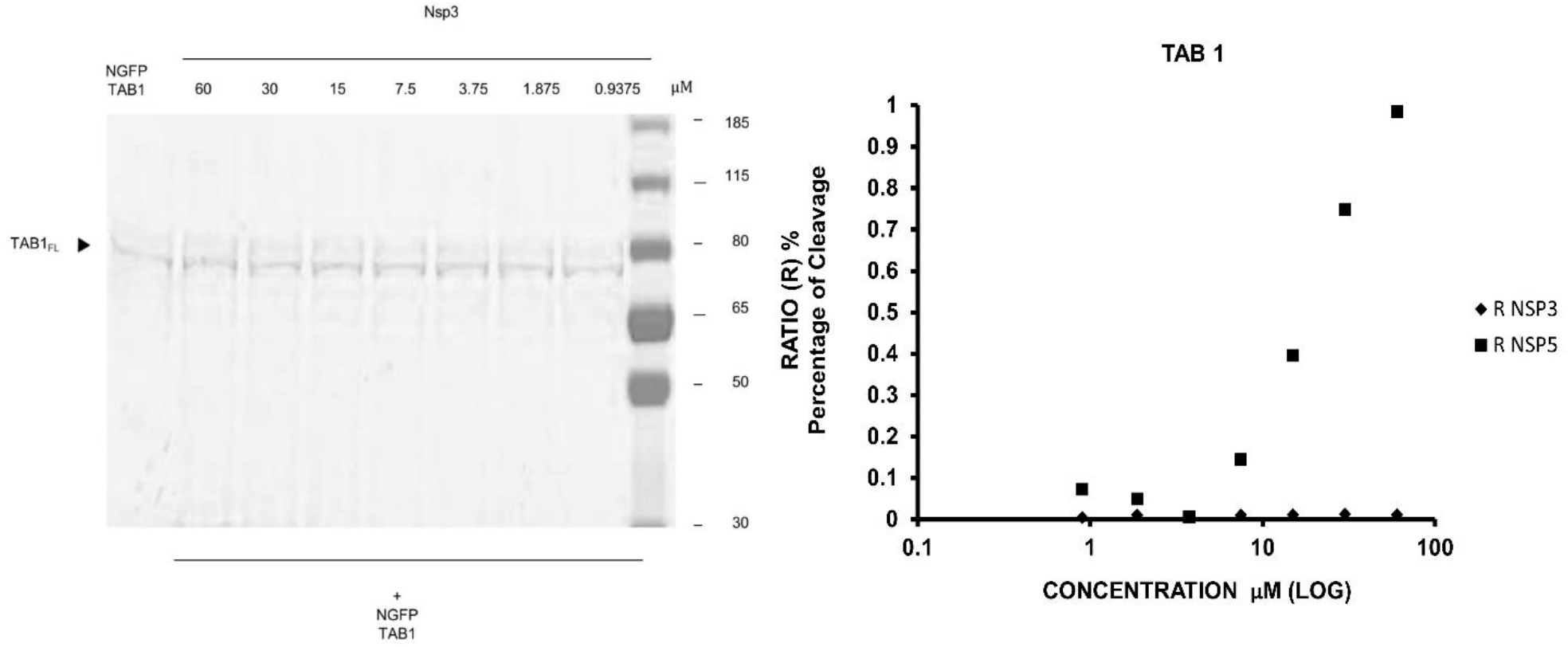
quantification of TAB1 cleavage with 3CLpro and PLpro. (left): SDS-page analysis of the cleavage of TAB1 protein, with a N-terminal GFP tag. The protein was expressed alone or in the presence of increasing concentrations of the SARS-CoV-2 protease PLpro (nsp3). The gel shows no cleavage site. (right): Dose-response curve of SARS-CoV-2 protease PLpro (nsp3) and 3CLpro9 (nsp5) on Human TAB1. The result from the SDS-page gel was processed using IMAGE J and the Percentage of cleavage (Ratio R) was calculated by measuring the length of the line, in pixels.

**Supplementary Figure 7:**
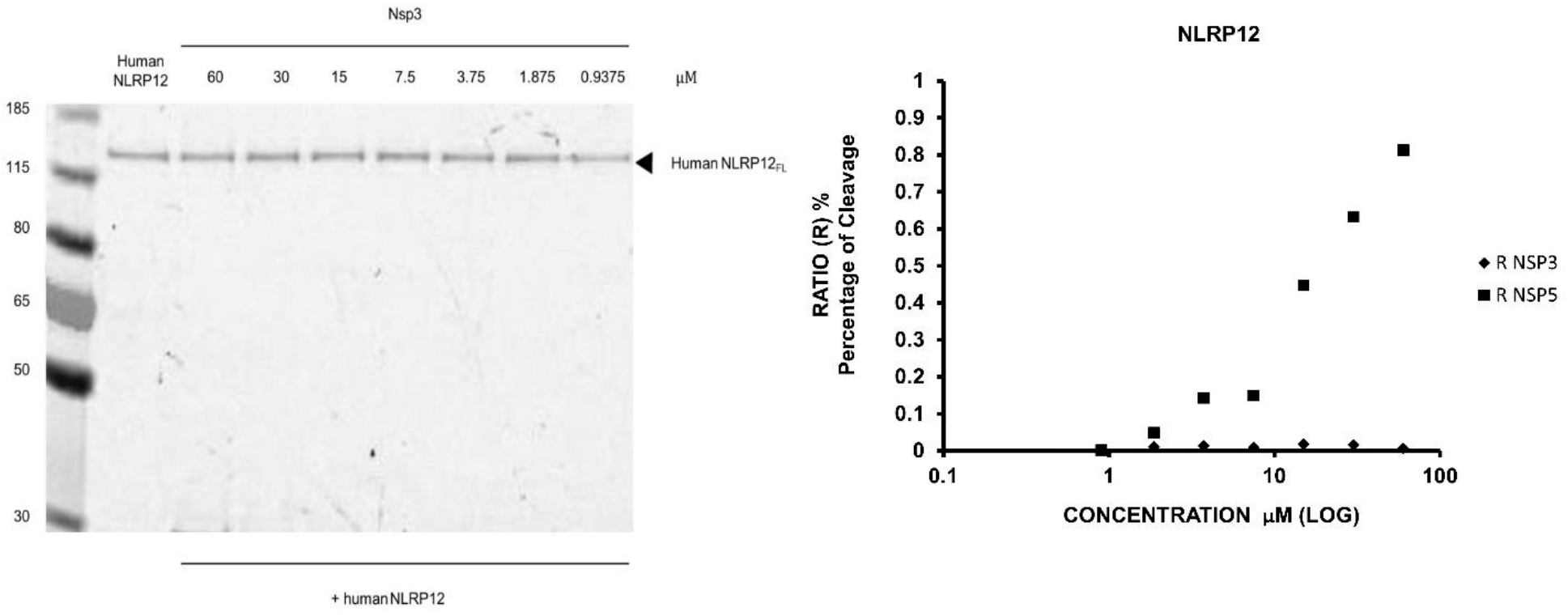
quantification of NLRP12 cleavage with 3CLpro and PLpro. (left): SDS-page analysis of the cleavage of NLRP12 protein, with a N-terminal Cherry tag. The protein was expressed alone or in the presence of increasing concentrations of the SARS-CoV-2 protease 3CLpro9 (nsp5). The gel shows no cleavage site. (right): Dose-response curve of SARS-CoV-2 protease PLpro (nsp3) and 3CLpro9 (nsp5) on Human NLRP12. The result from the SDS-page gel was processed using IMAGE J software and the Percentage of cleavage (Ratio R) was calculated by measuring the length of the line, in pixels.

**Supplementary Figure 8:**
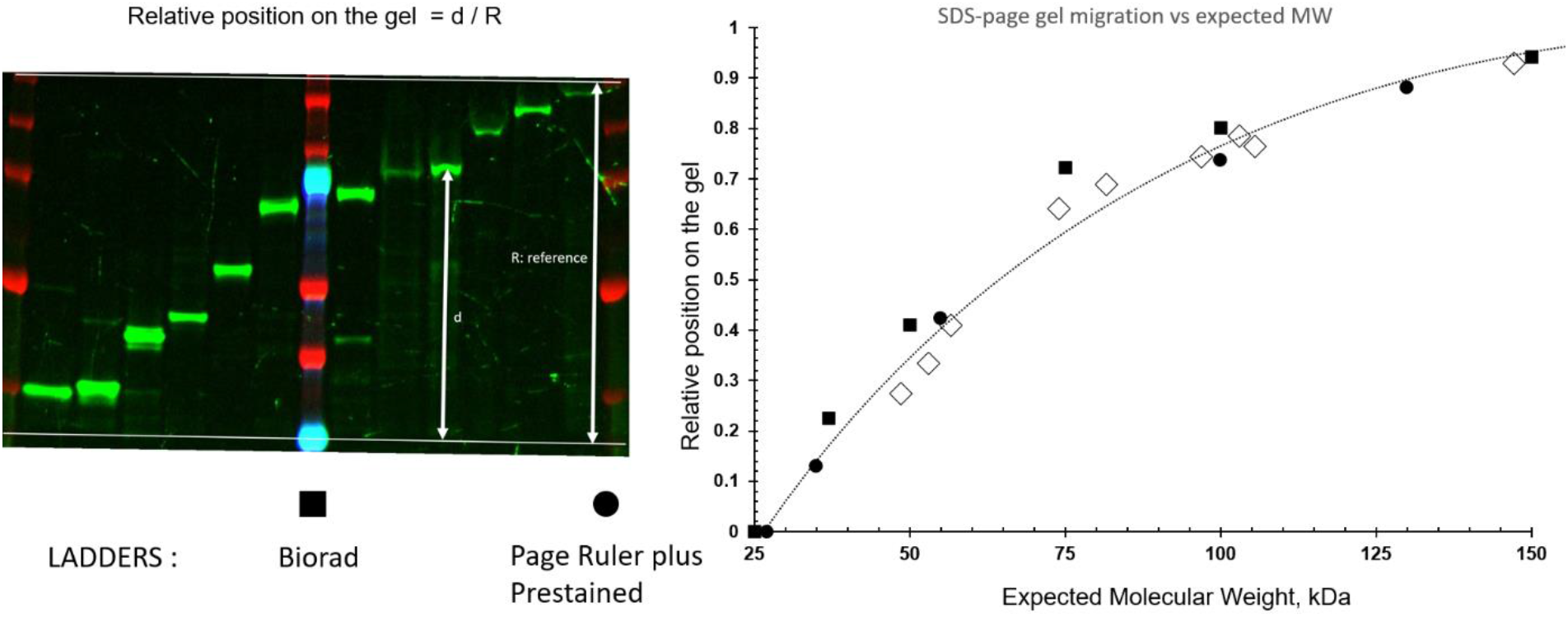
calibration of the migration of GFP-tagged proteins on SDS-page. **(left):** a range of HIIPs were selected with different molecular weights, ranging from 15 to 140 kDa; the HIIPS were expressed in LTE, mixed with LDS after 3h of expression, and separated on the SDS-page gel. In this gel, two different ladders were used to calibrate migration of 12 proteins of interest. The Page ruler Prestained ladder has been developed specifically for the 4-12% Bis-Tris gels (ThermoFisher, black ●) and seems more accurate to predict sizes of GFP-tagged proteins (in white diamonds ◊). The trendline indicated on the graph (right) will be used to estimate the size of the GFP-tagged fragments of TAB1 and NLRP12.

**Supplementary Figure 9:**
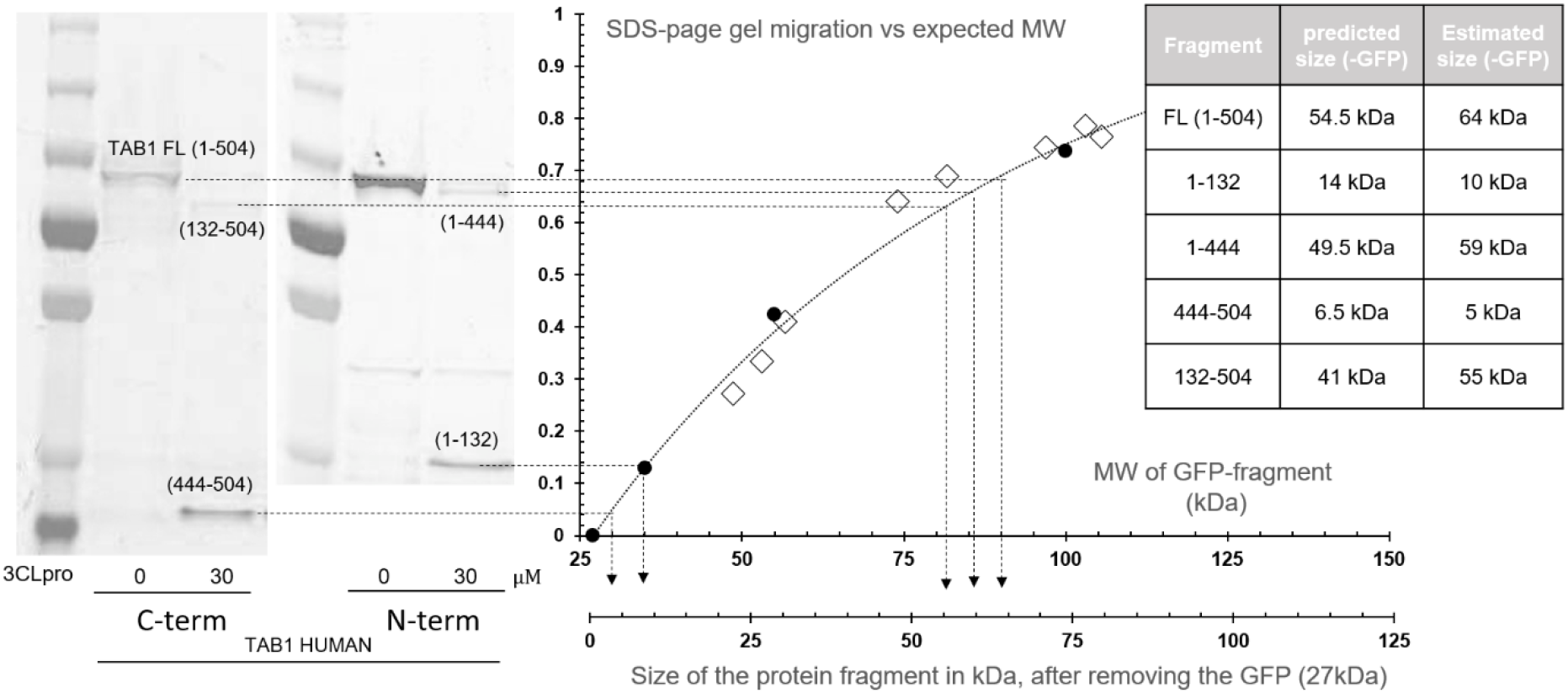
analysis of TAB1 fragments upon 3CLpro cleavage. Probably due to the non-denaturation of the protein TAB1 in our LDS loading protocol, the full-length protein human TAB1 migrates slower than expected on the SDS-page gel (estimated size 64 kDa from the calibration, vs 54.4 kDa expected). Nevertheless, the variations in size due to 3CLpro cleavage are consistent with the two sites at position 132 and 444. On the gel, the fragments are indicated, and the migration was analysed on the master curve to estimate the size of the GFP-tagged fragments. The 27kDa of the GFP were taken into account in the table (right) to compare the predicted and observed sizes of the proteins and protein fragments.

**Supplementary Figure 10:**
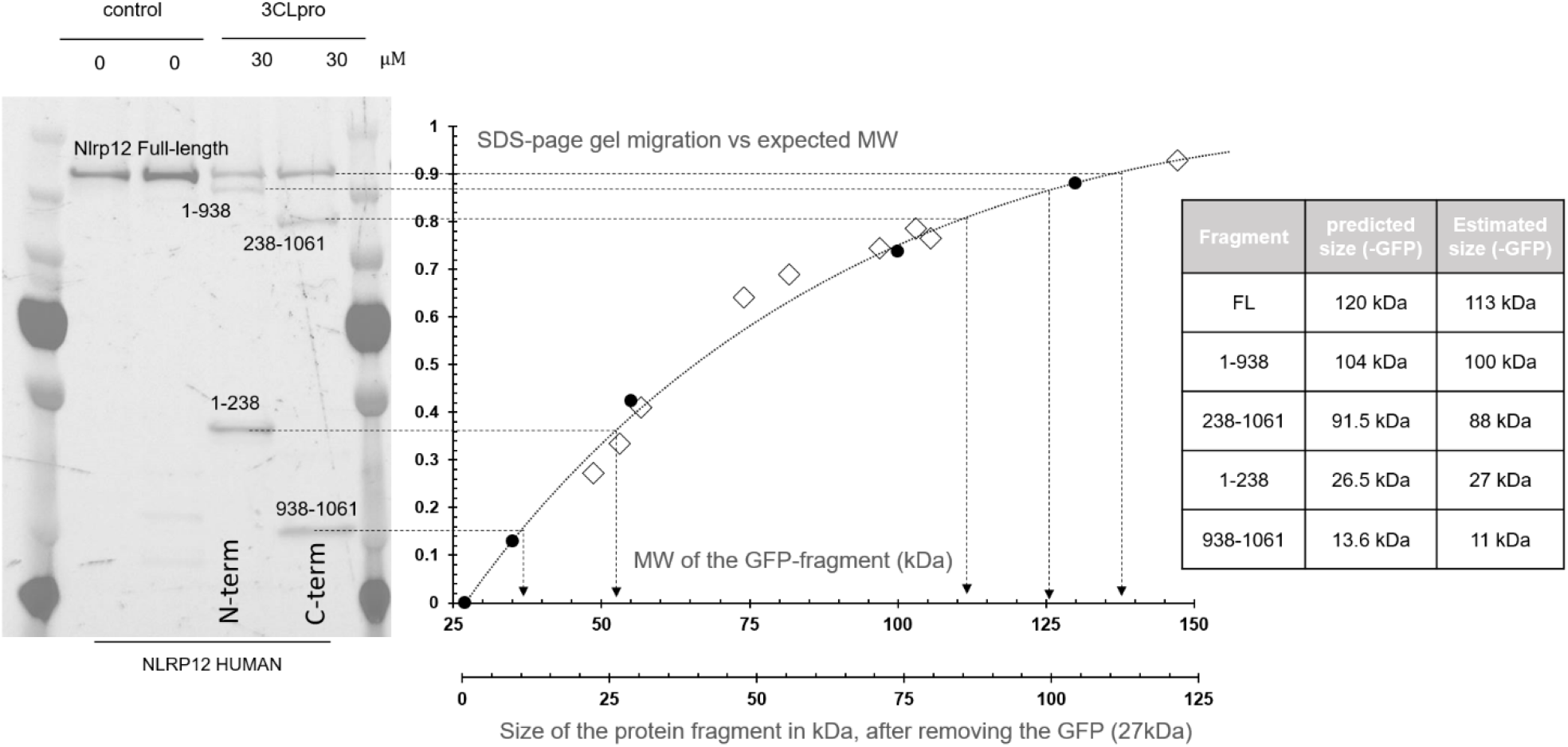
analysis of NLRP12 fragments upon 3CLpro cleavage. The full-length human NLRP12 proteins, tagged in N-term and C-term with GFP, were mixed with 3CLpro and analysed by SDS-page gels. The five bands obtained for the full-length NLRP12 and the four fluorescent cleavage products were analysed on the predictive size/migration plot. The variations in size are perfectly consistent with the two cleavages sites at position 238 and 938. On the SDS-page gel on the left, the fragments are indicated, and the migration was analysed on the master curve to estimate the size of the GFP-tagged fragments. The sizes obtained (after removing the 27kDa contribution of the GFP) were compared to the predicted sizes of the proteins and protein fragments.

